# YY1 Enhances the Stability of HIF-1α Protein by Interacting with NUSAP1 in Macrophages within the Prostate Cancer Microenvironment

**DOI:** 10.1101/2024.11.07.622424

**Authors:** Wenchao Li, SaiSai Chen, Jian Lu, Weipu Mao, Shiya Zheng, Minhao Zhang, Tiange Wu, Yurui Chen, Kai Lu, Chunyan Chu, Chuanjun Shu, Yue Hou, Xue Yang, Naipeng Shi, Zhijun Chen, Lihua Zhang, Lei Zhang, Rong Na, Ming Chen, Shenghong Ju, Dingxiao Zhang, Yi Ma, Bin Xu

## Abstract

Immune checkpoint therapy for prostate cancer (PCa) has failed in clinical trials; however, the precise underlying mechanisms involved remain elusive. PCa, a classic "immune-cold” tumor, is characterized by an immunosuppressive tumor microenvironment. Within this milieu, macrophages, the predominant immune cell population, have a propensity to infiltrate the hypoxic zones of tumors. In a previous study, we showed that Yin Yang 1 (YY1) is highly expressed in macrophages in PCa tissues. Here, through multiplexed imaging mass cytometry (IMC) of a PCa tissue microarray, we further demonstrate that YY1^+^ macrophages aggregate in hypoxic areas of tumors and that hypoxia promotes the phase separation of YY1 in the nucleus by increasing YY1 tyrosine phosphorylation in macrophages. Furthermore, YY1 binds to NUSAP1 and promotes the SUMOylation of HIF-1α, which promotes phase separation and stabilization of the HIF-1α protein. We also demonstrated that either treatment with a small molecule inhibitor (tenapanor) to decrease the YY1–NUSAP1–HIF-1α interaction or myeloid-specific YY1 gene knockout impairs subcutaneous PCa tumor formation. Furthermore, we present a first-generation tetrahedral DNA nanostructure (TDN) based on the proteolysis targeting chimera (PROTAC) technique, named YY1-DcTAC, which targets and degrades YY1 in tumor-associated macrophages. In a PCa mouse model, YY1-DcTAC exhibited prolonged drug efficacy, robust macrophage-specific responsiveness, potent antitumor effects, and increased CD8^+^ T cell tumor infiltration. In summary, our findings underscore the pivotal role of YY1 within the hypoxia/HIF-1α pathway in tumor-associated macrophages and affirm the therapeutic potential of targeting YY1 for treating PCa.

## INTRODUCTION

Prostate cancer (PCa) is one of the most prevalent malignancies affecting men globally^1^. Despite the ability of androgen deprivation therapy to significantly impede disease progression, a considerable number of patients still experience metastasis or develop castration resistance, ultimately resulting in fatalities^2^. Immune checkpoint therapy (ICT) has demonstrated promising and durable antitumor effects in late-stage patients with certain tumors^3^. ICT promotes T-cell-mediated antitumor immunity by reactivating endogenous T cells "braked" by immune checkpoint molecules. However, PCa has been recognized as an "immune-cold" tumor that exhibits immunosuppressive characteristics and sparse T-cell infiltration^4, 5^. Clinical trials in PCa patients have shown poor responsiveness to ICT treatment. These circumstances highlight the importance of investigating the PCa tumor microenvironment (TME) and identifying novel therapeutic targets.

Solid tumors are characterized by a hypoxic microenvironment^6^, which can promote malignant proliferation, invasion, and metastasis by modulating antitumor immunity^7, 8^. HIF-1α, a pivotal transcription factor with oxygen regulatory activity in the classic hypoxia-inducible factor (HIF) signaling pathway, regulates the tumor-specific immune response by promoting immune suppressive cell infiltration and activating multiple downstream signaling pathways and gene targets^9, 10^. Studies have highlighted the propensity of hypoxic environments to attract infiltrating macrophages^11^. Macrophages, the crucial component of the TME, have been established as a predominant immune cell subtype in most immune-cold tumors^12-14^. In various tumor types, including prostate cancer, breast cancer, and melanoma, infiltration of tumor-associated macrophages (TAMs) is significantly correlated with poor prognosis and tumor progression^15-17^. Mechanistically, TAMs can suppress the antitumor T-cell response by inducing tissue fibrosis, producing multiple immunosuppressive cytokines and chemokines, and decreasing the tumoral infiltration of CD8^+^ T cells^18, 19^.

In the present study, we demonstrate that TAMs with high Yin Yang 1 (YY1) expression aggregate in PCa hypoxic zones and that hypoxia induces phase separation of YY1 within the nuclei of macrophages. This process involves the interaction between YY1 and nucleolar and spindle-associated protein 1 (NUSAP1), which promotes the SUMOylation of HIF-1α, leading to increased phase separation and stabilization of HIF-1α. Moreover, we showed that treating PCa tumors with the small molecule inhibitor tenapanor, which disrupts the YY1–NUSAP1–HIF-1α interaction, effectively inhibits PCa tumor growth. Additionally, using a myeloid cell-specific YY1 knockout mouse model, we observed that YY1 deficiency in macrophages impairs subcutaneous tumor formation. To overcome the challenges posed by the structural heterogeneity and lack of active sites in YY1, which hinder drug development, we devised a tetrahedral DNA nanostructure based on the proteolysis-targeting chimera (PROTAC)^20^ technique to degrade YY1 in macrophages. We then validated this nanostructure’s pharmacokinetic characteristics and therapeutic efficacy in PCa. Overall, our findings provide novel insights into the role of YY1 and HIF-1α in TAMs, opening up novel potential therapeutic avenues for PCa patients.

## RESULTS

### YY1^+^ macrophages accumulate in hypoxic PCa tissues

Imaging mass cytometry (IMC) was used to analyze prostate tissue microarrays comprising 46 prostate cancer (PCa) tissues and 26 adjacent noncancerous tissues (**Figure 1A**). A comprehensive panel of markers was used for analysis, including the hypoxia marker HIF-1α, the macrophage marker CD68, the transcription factor YY1, and the cell nuclear marker DNA1. A discernible cellular landscape within PCa tissues emerged amidst the intricate spatial orchestration of cell subtypes and transcription factor expression profiles, particularly within hypoxic regions marked by HIF-1α. Notably, a significantly greater percentage of YY1^+^ macrophages than YY1^-^ macrophages was detected within these hypoxic regions (**Figure 1B**). Interestingly, no substantial differences in infiltration were observed between YY1^+^ or YY1^-^ macrophages within the hypoxic zones of adjacent noncancerous tissues (**Figure 1C**), suggesting a context-specific role for YY1 in the tumor setting.

**Figure 1.**
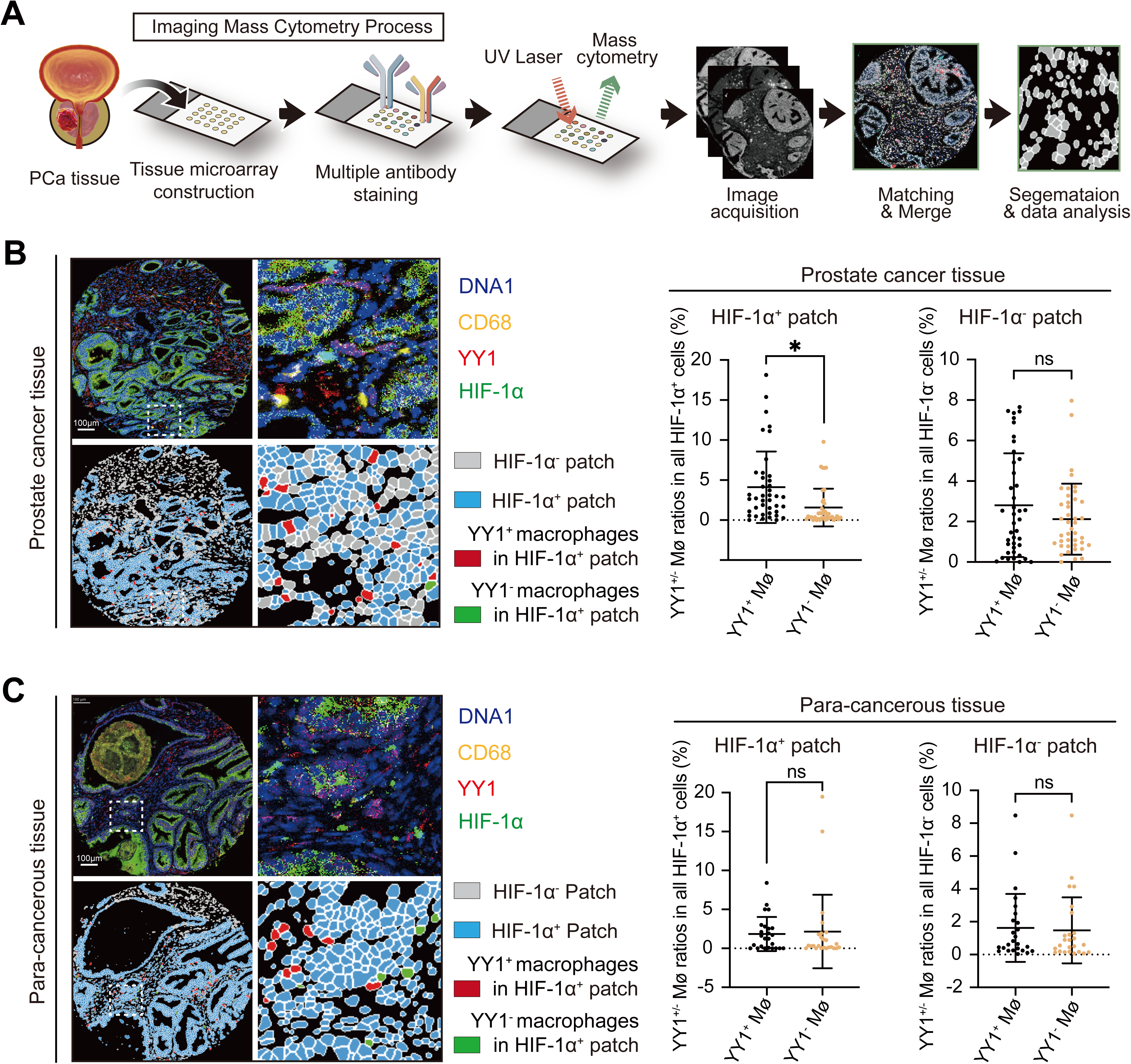
YY1^+^ macrophages accumulate in hypoxic tumor tissues. **(A)** Schematic diagram showing the process of imaging mass cytometry (IMC) using prostate tissue. **(B–C)** IMC of the indicated markers in prostate cancer (PCa) patient tumor and para-cancerous tissues. Dot plots showing the ratio of YY1^+^ or YY1^-^ macrophages in HIF-1α^+^ patches. \**P* < 0.05.

Moreover, our investigation extended to areas lacking HIF-1α expression (HIF-1α-negative patch), where the positivity rate of YY1 in macrophages within nonhypoxic regions exhibited no significant difference, whether in the tumor center or adjacent noncancerous tissues (**Figures 1B–C**). Overall, the infiltration of YY1-expressing macrophages in hypoxic PCa tissues highlights their potential relevance in responding to hypoxia.

### Hypoxia enhances YY1 phase separation by inducing tyrosine phosphorylation of YY1 in macrophages

To further investigate the response of macrophages to hypoxia, we utilized cobalt chloride (CoCl_2_) to induce hypoxia *in vitro* in THP-1 cells. Interestingly, we observed the accumulation of punctate YY1 particles within the cell nucleus under hypoxic conditions, which could be attenuated by the phase separation inhibitor 1,6-hexanediol (1,6-Hex) (**Figure 2A**). The formation of phase-separated droplets has been shown to participate in the cellular response to physical stimuli or stress and typically relies on the presence of intrinsically disordered protein regions (IDRs)^21^. Here, we predicted through the Predictor of Natural Disordered Regions (PONDR) website that YY1 contains a lengthy N-terminal IDR segment (**Figure S1A**). Confocal microscopy analysis of living cells constructed with the YY1-IDR-EGFP or YY1-non-IDR-EGFP plasmid revealed that the phase separation ability of YY1-IDR-EGFP, but not that of YY1-non-IDR-EGFP, increased under hypoxic conditions (**Figure S1B**). Furthermore, we conducted a time-gradient experiment involving hypoxia and reoxygenation in THP-1 cells. As the duration of hypoxia increased to 8 h, the droplet density gradually increased, returning to baseline levels after 120 min of reoxygenation (**Figure 2B**).

**Figure 2.**
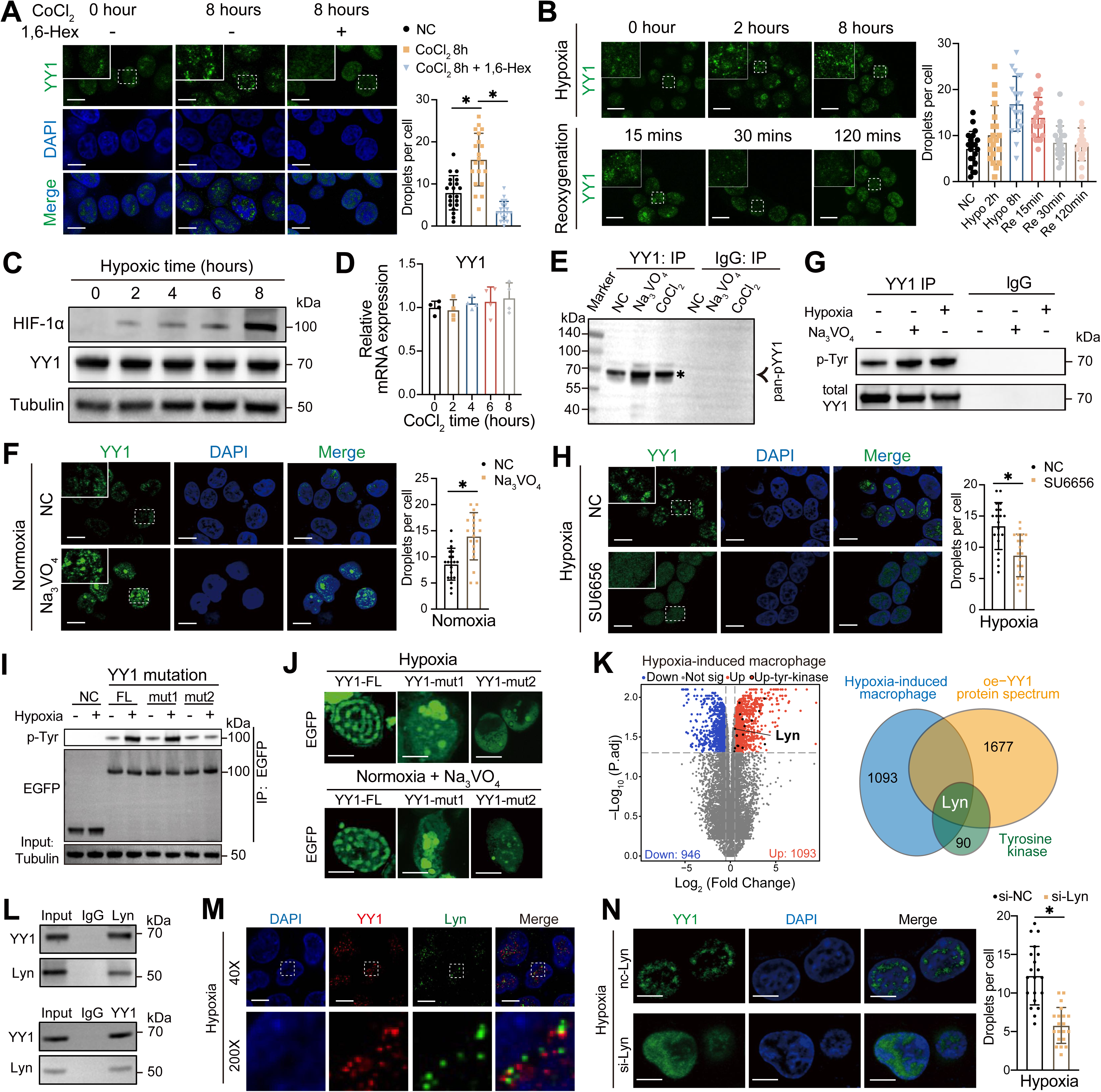
Hypoxia enhances YY1 phase separation by inducing YY1 tyrosine phosphorylation in macrophages. **(A)** Immunofluorescence staining of the indicated markers in THP-1 cells subjected to hypoxia and/or 1,6-hexanediol. **(B)** The localization of YY1 observed by immunofluorescence after hypoxia or reoxygenation for the indicated periods. **(C)** Western blot of HIF-1α and YY1 in THP-1 cells under different periods of hypoxia. **(D)** qRTLPCR analysis showing the relative RNA expression of *HIF-1*α and *YY1* in THP-1 cells under different periods of hypoxia. **(E)** Western blot analysis using an anti-phosphorylated YY1 antibody to detect YY1 or IgG immunoprecipitated proteins under hypoxia or Na_3_VO_4_ treatment. **(F)** Immunofluorescence staining of the indicated markers in THP-1 cells treated with Na_3_VO_4_ or control solution under normal oxygen conditions. **(G)** Western blot analysis using a tyrosine phosphorylation antibody to detect YY1 or an IgG immunoprecipitation protein under hypoxia or Na_3_VO_4_ treatment. **(H)** Immunofluorescence staining of the indicated markers in THP-1 cells treated with SU6656 or control solution under hypoxia. **(I)** Western blot analysis using a tyrosine phosphorylation antibody to detect EGFP immunoprecipitated from YY1 full-length (YY1-FL), YY1 mutation 1 (YY1-mut1, including amino acid sites 8 and 383) and YY1 mutation 2 (YY1-mut2, including amino acids 145, 185, 251, and 254)-transfected THP-1 cells. **(J)** Immunofluorescence of YY1-FL-, YY1-mut1- and YY1-mut2-transfected THP-1 cells. **(K)** The left volcano plot shows proteins demonstrating upregulated (red) and downregulated (blue) expression in the spectra from THP-1-induced macrophages subjected to hypoxia compared to normal control macrophages, and the black-circled red points indicate tyrosine kinases demonstrating upregulated expression. The right schematic panel shows the overlap indicated to identify Lyn. **(L–M)** Co-IP and immunofluorescence of YY1 and Lyn from hypoxia-induced THP-1 cells. N, Immunofluorescence of the indicated markers in THP-1 cells transfected with si-Lyn or the normal control under hypoxia. All the immunofluorescence data were analyzed, and twenty cells randomly selected from three repeated groups were counted. Scale bar, 5 μm. * *P* < 0.05.

To investigate the mechanisms underlying the abovementioned phenotypes, we evaluated the expression of YY1 under hypoxia and found that the expression level of YY1 was not altered by varying the concentrations of CoCl_2_ or durations of exposure (**Figures 2C, D,** and **S1C**). Accumulating evidence suggests that the phosphorylation of proteins enhances their phase separation^22, 23^; thus, we further investigated the impact of hypoxia on YY1 protein phosphorylation. First, using a panphosphorylated antibody, we revealed that hypoxia augments the expression of phosphorylated YY1 (**Figure 2E**). Sodium orthovanadate (Na_3_VO_4_), a commonly used protein phosphatase inhibitor, not only enhanced the expression of panphosphorylated YY1 (**Figure 2E**) but also facilitated the formation of phase separation droplets of YY1 (**Figure 2F**) under normoxia. Serine and tyrosine are frequently reported phosphorylation sites of YY1 (p-Ser and p-Tyr), and we observed that YY1 p-Tyr but not p-Ser levels significantly increased under hypoxia (**Figures 2G** and **S1D**). In previous studies, Src family kinases were revealed to be important mediators of YY p-Tyr ^24^. In this study, our results revealed that SU6656, a selective Src family kinase inhibitor, mitigated YY1 hypoxia-induced phosphorylation and phase separation (**Figures 2H** and **S1E**). These findings suggest that hypoxia augments YY1-mediated phase separation by promoting YY1 tyrosine phosphorylation.

To elucidate the specific sites of hypoxia-induced YY1 p-Tyr, we strategically engineered six tyrosine residues within YY1 into two distinct mutant groups, each fused with EGFP: YY1-mut1 (sites Y8F and Y383F) and YY1-mut2 (sites Y145F, Y185F, Y251F, and Y254F), based on their respective locations within the YY1-non-IDR or YY1-IDR domains (**Figure S1A**). Interestingly, YY1-mut2 (mutations at the p-Tyr site within the YY1-IDR region) but not YY1-mut1 (mutations in the YY1-non-IDR domain) effectively inhibited the hypoxia-induced increase in the p-Tyr level (**Figure 2I**). Morphological assessments also revealed that YY1-mut2 effectively mitigated the enhancement of phase separation induced by Na_3_VO_4_ or hypoxia (**Figure 2J**).

Furthermore, to determine the tyrosine kinases responsible for the hypoxia-induced phosphorylation of YY1, we compared the mass spectra of hypoxia-induced genes identified via macrophage RNA-seq and YY1 pull-down protein analysis. One Src family kinase, Lyn, was subsequently identified. Subsequently, through protein immunoprecipitation and cellular immunofluorescence assays, we confirmed the interaction between YY1 and Lyn and their colocalization in hypoxia-treated THP-1 cells (**Figures 2L, M**). In addition, silencing Lyn via small interfering RNA markedly impeded the formation of YY1 droplets under hypoxic conditions (**Figures 2N** and **S1F**). Briefly, our investigation revealed that hypoxia promotes the aggregation of YY1 into nuclear puncta by inducing YY1 phosphorylation, with Lyn emerging as a pivotal kinase mediating this process.

Moreover, the YY1 C-terminal segment (comprising amino acids 296–414) constitutes a highly conserved DNA-binding region that houses four zinc-finger motifs (amino acids 296–320, 325–347, 353–377, and 383–407)^25^. In our investigation, we found that silencing YY1 amino acids 296–320 (YY1^△296-320^-EGFP), a zinc-finger motif located within the functional YY1-IDR region, significantly diminished the formation of phase separation droplets under hypoxia. Conversely, the general silencing of the other three motifs (YY1^△321-414^-EGFP) did not have a notable impact on the formation of phase-separated droplets (**Figure S1G**). Collectively, our findings underscore the critical roles of the p-Tyr sites (145, 185, 251, 254) and DNA-binding regions (amino acids 296–320) within the YY1-IDR domain in mediating hypoxia-induced YY1 phase separation.

### YY1 stabilizes HIF-1α by inhibiting its ubiquitination

To further investigate the mechanistic processes underlying the function of YY1 in macrophages under hypoxic conditions, we conducted RNA sequencing analyses of macrophages overexpressing YY1 and those subjected to hypoxia. The results showed a substantial overlap in the enrichment pathways between these two groups of macrophages (**Figure 3A**). Notably, among the top ten pathways enriched in both treatment groups, the HIF-1 signaling pathway was strongly enriched (**Figures 3B, C**). Given that HIF-1α plays a fundamental role in the hypoxia signaling pathway^26, 27^, we used siRNA to inhibit YY1 expression and observed its impact on HIF-1α expression in THP-1 cells. Western blotting revealed a notable decrease in HIF-1α protein levels in THP-1 cells transfected with siYY1 compared to those in control cells (**Figure 3D**), while the mRNA levels of HIF-1α remained unaffected (**Figure 3E**). This finding suggested the involvement of a posttranslational mechanism that potentially stabilizes HIF-1α via YY1 in macrophages. Consistent with this hypothesis, the cycloheximide (CHX) chase assay demonstrated a significant reduction in the stability of HIF-1α upon YY1 suppression (**Figure 3F**). In addition, we observed that the proteasome inhibitor MG132 increased HIF-1α expression during hypoxia and counteracted the inhibitory effect of YY1 knockdown on HIF-1α levels (**Figure 3G**). The HIF-1α protein is known to be stabilized under hypoxic conditions and degraded through the ubiquitination-proteasome pathway in a normoxic environment^28, 29^. Herein, we conducted a ubiquitin assay to assess the ubiquitination level of HIF-1α, revealing an increase in the ubiquitination level of HIF-1α following YY1 suppression (**Figure 3H**). Moreover, silencing YY1 in THP-1 cells led to increased degradation of HIF-1α, which was mediated by descending small ubiquitin-related modifier 2/3 (SUMO 2/3)-induced SUMOylation (**Figure 3I**). In summary, these findings confirm that YY1 promotes the stabilization of the HIF-1α protein, a process closely linked to its ubiquitination and SUMOylation.

**Figure 3.**
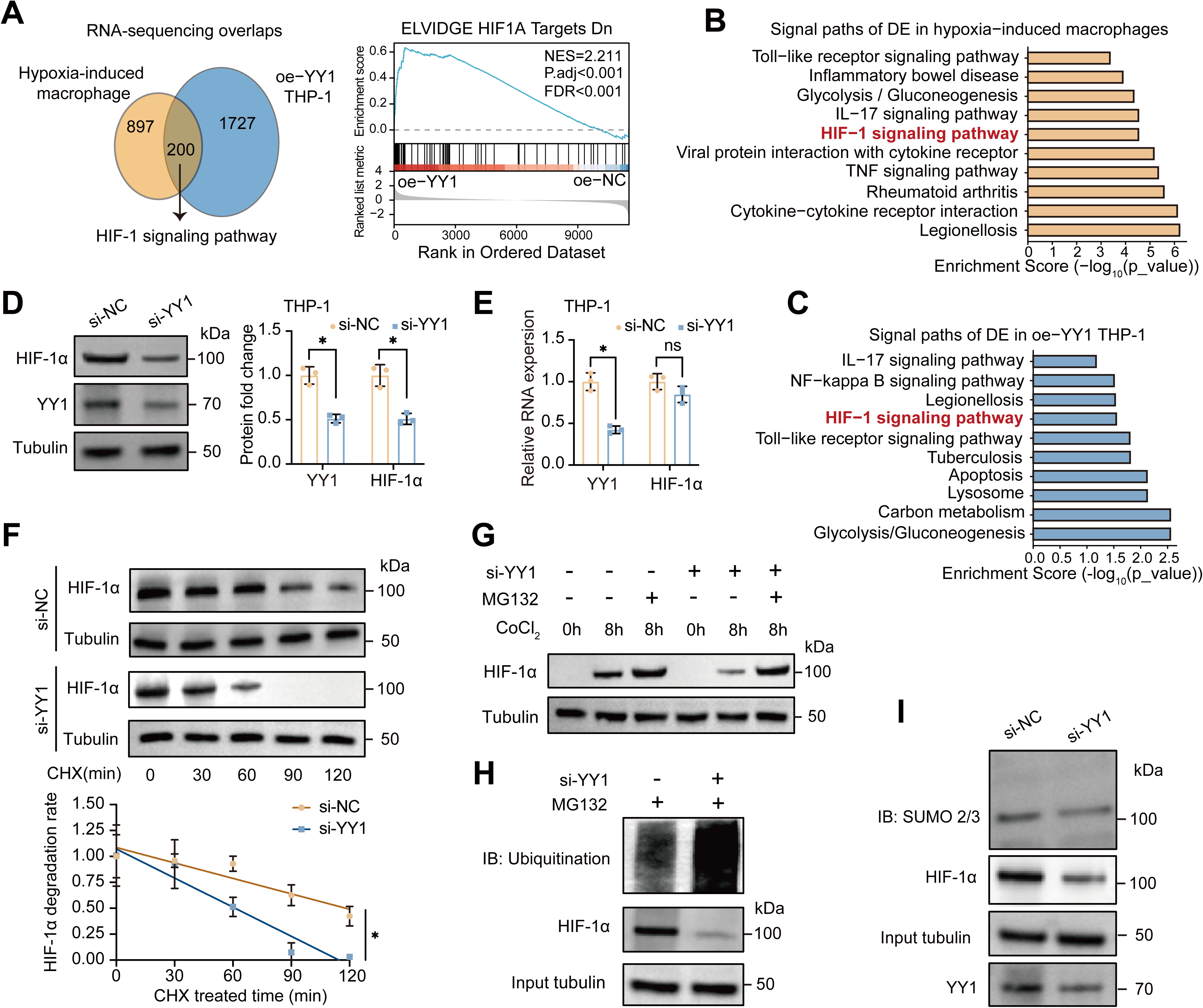
YY1 stabilizes HIF-1α by promoting its SUMOylation and inhibiting its ubiquitination. **(A–C)** GSEA revealed that the HIF-1 signaling pathway was upregulated in THP-1 cells subjected to hypoxia or transfected with oe-YY1 lentivirus. **(D)** Western blot analysis showing the expression of HIF-1α and YY1 in hypoxia-induced THP-1 cells transfected with siYY1 or the normal control. **(E)** qRTLPCR showing the expression of YY1 and HIF-1α in THP-1 cells transfected with si-YY1 or the normal control. **(F)** Western blot analysis of HIF-1α expression in hypoxia-induced THP-1 cells treated with cycloheximide (CHX) for different durations. **(G)** Western blot analysis of HIF-1α expression in THP-1 cells treated with MG132. **(H)** Ubiquitin experiments showed that the ubiquitination of HIF-1α was enhanced after YY1 was suppressed in hypoxia-induced THP-1 cells. **(I)** SUMO 2/3-mediated SUMOylation in si-YY1- or control-transfected THP-1 cells under hypoxia. * *P* < 0.05.

### NUSAP1 promotes HIF-1α SUMOylation and inhibits its ubiquitination

To further explore the mechanism by which YY1 stabilizes HIF-1α in macrophages, we performed mass spectrometry analysis of the YY1 protein in THP-1 cells. The results revealed NUSAP1 as a protein bound by YY1, and co-IP experiments confirmed the direct interaction between YY1 and NUSAP1 (**Figures 4A, B**). Elevated expression of NUSAP1, a microtubule-associated protein, in the TME is correlated with tumor infiltration of myeloid-derived immune cells^30^. Notably, NUSAP1 harbors an SAP domain commonly found in small ubiquitin-related modifier (SUMO) ligases that is associated with substrate recognition and ligase activity^31^. Given the role of YY1 in stabilizing HIF-1α in macrophages, we extended our investigation to ascertain whether NUSAP1 contributes to HIF-1α expression. Remarkably, silencing NUSAP1 led to substantial downregulation of HIF-1α expression (**Figure S2A**), and the MG132–CHX assay showed that NUSAP1 could bolster HIF-1α stabilization (**Figures 4C** and **S2B**). Consistent with the stabilizing effect of YY1 on HIF-1α, the ubiquitination of HIF-1α markedly decreased after NUSAP1 knockdown (**Figure 4D**).

**Figure 4.**
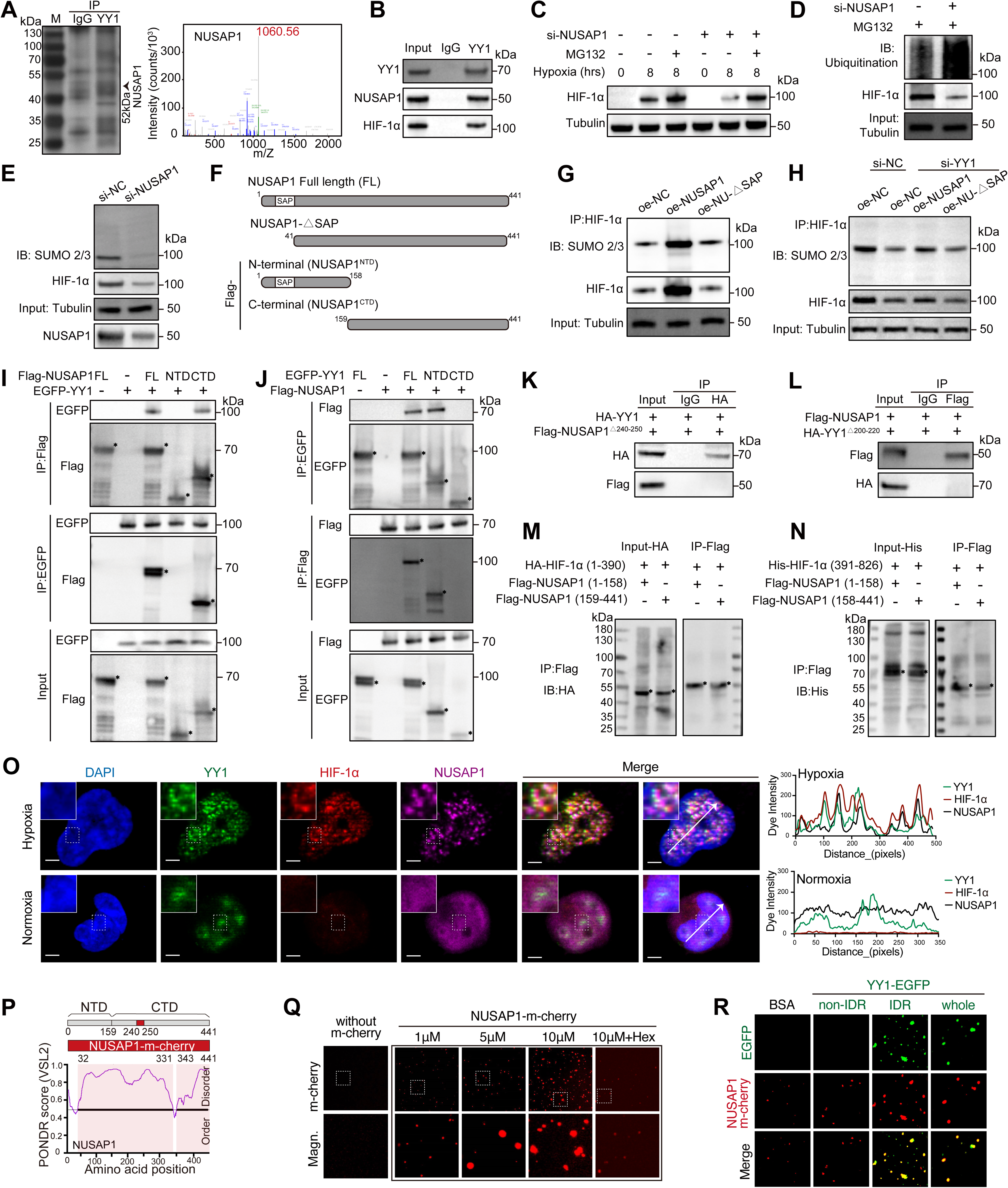
YY1 promotes the stability of HIF-1α by binding to NUSAP1. **(A)** Mass spectrometry of proteins immunoprecipitated with YY1 and IgG. **(B)** Co-IP of proteins immunoprecipitated with YY1 and IgG showed binding between YY1 and NUSAP1 in hypoxia-induced THP-1 cells. **(C)** Western blot analysis of HIF-1α expression in THP-1 cells treated with MG132. **(D)** Ubiquitin experiments showed that the ubiquitination of HIF-1α was enhanced after NUSAP1 was suppressed in hypoxia-induced THP-1 cells. **(E)** Western blot analysis showing SUMO 2/3-mediated SUMOylation in si-NUSAP1- or si-NC-transfected THP-1 cells. **(F)** Diagram of the SAP domain and N/C-terminus of NUSAP1. **(G–H)** SUMOylation assays in hypoxia-induced THP-1 cells under the indicated conditions. **(I)** The interaction of EGFP-YY1 with truncated NUSAP1 in hypoxia-induced THP-1 cells. **(J)** The interaction of Flag-NUSAP1 with truncated YY1 in hypoxia-induced THP-1 cells. **(K–L)** Co-IP assays showing the interactions of the indicated HA-YY1 or Flag-NUSAP1 mutant. **(M–N)** Co-IP assays showing the interactions of truncated HIF-1α and NUSAP1 proteins in hypoxia-induced THP-1 cells. **(O)** Immunofluorescence assays showing the colocalization of the indicated markers. The dye intensity alongside the white lines was calculated and plotted. Scale bar, 2 μm. **(P)** Schematic diagram of the NUSAP1 IDR. **(Q)** Fluorescence images of m-cherry showing droplet formation under different concentrations of NUSAP1-m-cherry protein. **(R)** Fluorescence images of EGFP and m-cherry showing the physical colocalization of NUSAP1-m-cherry with the YY1-IDR-EGFP or YY1-non-IDR-EGFP.

SUMOylation has been shown to counteract the ubiquitination of substrates under specific circumstances ^32, 33^. Herein, we found that knocking down NUSAP1 reduced the SUMOylation level of HIF-1α (**Figure 4E**). Furthermore, to elucidate the specific role of the SAP domain in this regulatory process, we generated a NUSAP1-△SAP variant lacking the SAP domain and assessed its impact on SUMOylation and HIF-1α ubiquitination (**Figure 4F**). Notably, while full-length NUSAP1 promoted HIF-1α SUMOylation, the NUSAP1-△SAP variant failed to elicit similar effects (**Figure 4G**). Moreover, our investigations revealed the involvement of YY1 in mediating the SUMOylation of HIF-1α by NUSAP1. Knockdown of YY1 decreased the SUMOylation of HIF-1α facilitated by NUSAP1, while overexpression of NUSAP1 reversed HIF-1α SUMOylation and caused corresponding changes in HIF-1α protein levels (**Figure 4H**). These findings suggest that the interaction between YY1 and NUSAP1 facilitates the SAP domain-mediated SUMOylation of HIF-1α while inhibiting its ubiquitination, ultimately culminating in increased HIF-1α expression.

To validate the interaction between YY1 and NUSAP1, we predicted the binding sites by bioinformatics analysis, ranging mainly within amino acids 240–250 of NUSAP1 and amino acids 200–220 of YY1 (**Figure S2C**). In addition to the zinc finger structure domain (C-terminal DNA binding domain) mentioned above, the N-terminal domain (NTD) of YY1 has been reported to serve as the primary domain for protein modifications^34^. Here, we generated truncated forms of the N/C-terminal domains of YY1 (YY1^NTD^: 1–320, YY1^CTD^: 321–414) and NUSAP1 (NUSAP1^NTD^: 1–158, NUSAP1^CTD^: 159–441) and tagged them with Flag or EGFP, respectively (**Figures 4F** and **S2D**). Flag pull-down followed by co-IP assays confirmed the strong binding between YY1 and NUSAP1^CTD^ (Flag-NUSAP1^1–158^) but not between YY1 and NUSAP1^NTD^ (Flag-NUSAP1^159–441^) (**Figure 4I**). Similarly, YY1^NTD^ (EGFP-YY1^1–320^), but not YY1^CTD^ (EGFP-YY1^321–414^), was found to interact with NUSAP1 (**Figure 4J**). Next, we endeavored to precisely silence the regions harboring the most densely predicted sites within YY1^NTD^ and NUSAP1^CTD^, constructing truncated proteins tagged with Flag and HA, respectively. Our findings revealed a notable reduction in the binding between the YY1 and NUSAP1 proteins after silencing amino acids 240–250 of NUSAP1 and amino acids 200–220 of YY1. This finding suggests that the regions encompassing amino acids 240–250 of NUSAP1 and 200–220 of YY1 may represent the most crucial binding sites (**Figures 4K, L** and **S2E, F)**. Briefly, these results obtained from the truncated proteins validated the interaction between YY1 and NUSAP1. Concurrently, given the direct involvement of the SAP domain of NUSAP1 in mediating HIF-1α SUMOylation, we further examined the interaction between NUSAP1 and HIF-1α under hypoxic conditions. To discern specific binding regions, we subdivided HIF-1α into N-terminal (HA-HIF1α^1–390^) and C-terminal (His-HIF-1α^391–826^) fragments, each tagged accordingly. Through co-IP experiments, we confirmed the binding of both HIF-1α fragments to the NUSAP1 N/C-terminus (**Figures 4M, N**). In summary, our investigation of the truncated proteins mentioned above elucidate the interactions involving YY1, NUSAP1, and HIF-1α.

Moreover, we examined the morphological distribution of YY1, NUSAP1, and HIF-1α in cells under normoxic/hypoxic conditions via IF imaging to further investigate the associated intracellular processes. Our findings indicate that hypoxia triggers several notable phenomena. First, YY1 significantly increased the formation of liquid separation droplets under hypoxic conditions, consistent with previous observations. Second, the distribution of NUSAP1 in cells transitioned from a uniform dispersion to punctate aggregation. Third, the expression of HIF-1α increased under hypoxia, as expected, and HIF-1α formed droplets closely associated with or even colocalized with YY1 and NUSAP1 (**Figure 4O**). Surprisingly, almost the whole length of NUSAP1 was characterized as intrinsically disordered (**Figure 4P**), which has been recognized as a key element in phase separation. Then, we constructed an m-cherry-tagged NUSAP1 chimeric protein and conducted IF assays to examine its ability to form condensates with different protein concentrations in solution, in which PEG 8000 and 150 mM NaCl were used to mimic the crowding situation of the intracellular environment. As expected, significant aggregation of condensates was observed with increasing NUSAP1 chimera concentration, which could be reversed by 1,6-hex (**Figure 4Q**). Next, we mixed YY1-IDR and non-IDR chimeras tagged with EGFP with NUSAP1-m-cherry. Strikingly, we observed obvious colocalization of NUSAP1 with YY1-IDR and full-length YY1 (**Figure 4R**).

### HIF-1α undergoes SUMOylation-related phase separation under hypoxic conditions

Notably, recent studies have confirmed that SUMOylation of substrates and SUMO noncovalent interactions promote SUMOylation-mediated phase separation^35, 36^. To further clarify the role of phase separation in HIF-1α SUMOylation and stabilization, we generated EGFP-tagged HIF-1α chimeras based on the predicted HIF-1α IDRs, including IDR1-EGFP (EGFP-HIF-1α^1–80^), IDR2-EGFP (EGFP-HIF-1α^391–750^), and non-IDR-EGFP (EGFP-HIF-1α^81–390^) (**Figure 5A**). Fluorescence microscopic visualization of HIF-1α chimeric proteins revealed that EGFP-tagged IDR1 and IDR2 formed significantly enlarged condensates at concentrations ranging from 5–10 μM, which could be further inhibited by 1,6-hex, whereas non-IDR-EGFP did not yield noticeable condensates at 10 μM (**Figure 5B**). To replicate the behavior of IDRs within macrophages, we introduced segmented HIF-1α plasmids into THP-1 cells and conducted live imaging using confocal fluorescence microscopy without cell fixation. Under normoxic conditions, HIF-1α degraded rapidly, and no apparent condensates were observed. However, exposure to physical hypoxia led to a significant increase in fluorescence intensity, accompanied by the appearance of mobile condensates, confirming their liquid-like properties through fusion and fission processes (**Figure 5C**). Moreover, we conducted a fluorescence recovery after photobleaching (FRAP) assay in hypoxia-preconditioned THP-1 cells. The fluorescence intensity of the HIF-1α IDR1 protein almost completely recovered within 80 sec postbleaching, indicating that the condensed HIF-1α IDR1 underwent rapid exchange with the surrounding environment *in vivo* (**Figure 5D**). Subsequently, we subjected THP-1 cells transfected with EGFP-HIF-1α segmented plasmids to hypoxia to observe the cellular localization induced by hypoxia. Non-IDR chimeras exhibited a uniform pattern in the cytoplasm and nucleus, whereas IDRs formed clustered puncta mainly in the nucleus (**Figure 5E**). Furthermore, deletion of the IDR1 or IDR2 domains from the HIF-1α full-length plasmid suppressed phase-separated droplet formation. Notably, deletion of IDR1 resulted in predominant nuclear localization of the protein, whereas deletion of IDR2 led to an even distribution throughout the cells (**Figure 5F**), suggesting that the IDR2 domain may facilitate the nuclear entry of HIF-1α under hypoxic conditions.

**Figure 5.**
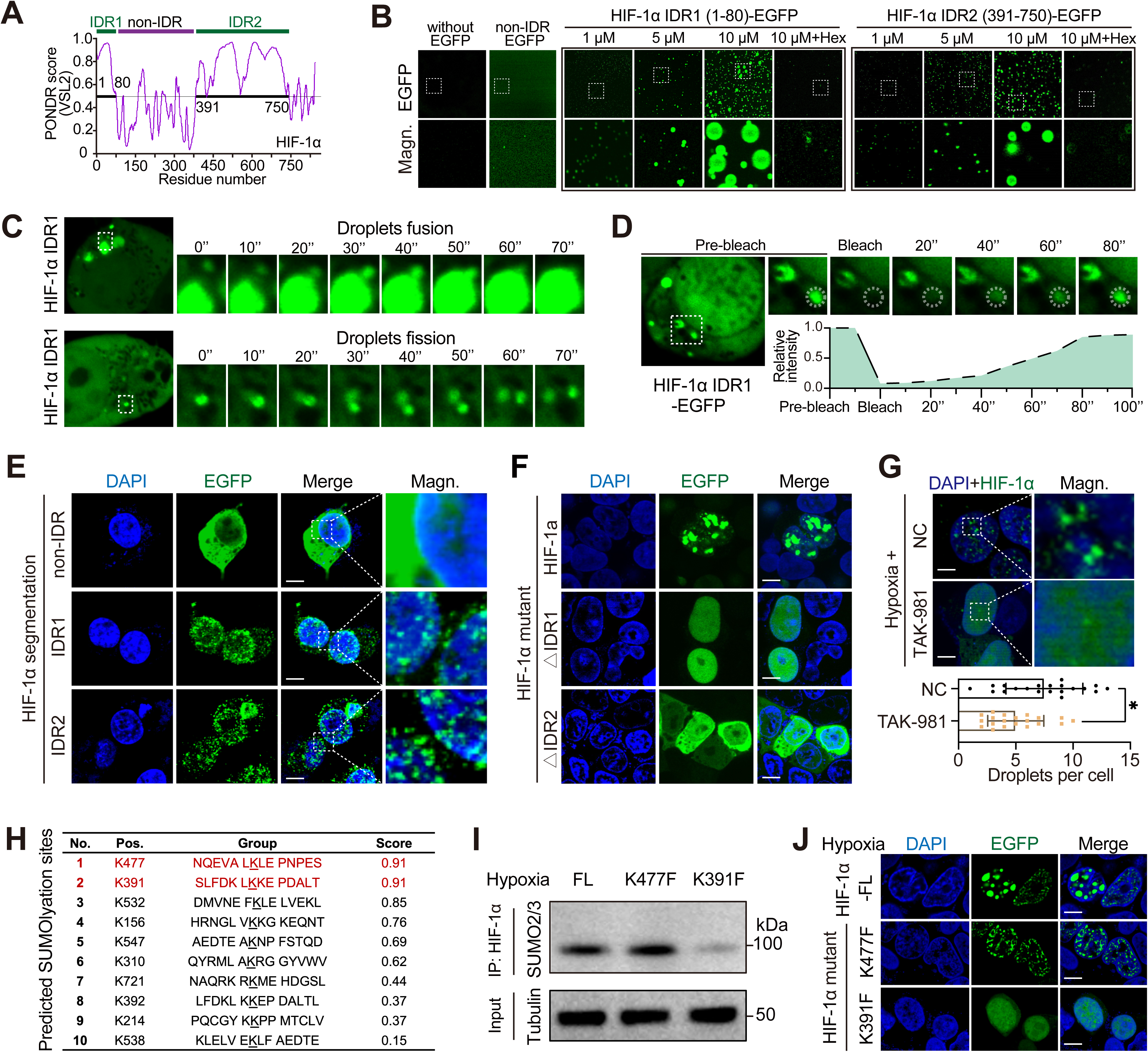
HIF-1α undergoes SUMOylation-related phase separation under hypoxia. **(A)** Diagram showing the intrinsically disordered protein regions (IDRs) of HIF-1α. **(B)** Representative images of phase separation condensates based on HIF-1α-IDR1/2-EGFP with different protein concentrations or with 1,6-hexanediol. **(C)** The process of fusion and fission in HIF-1α-IDR1-EGFP-mediated condensates. **(D)** Fluorescence intensity of the condensate during fluorescence recovery after photobleaching (FRAP) assay. The average relative fluorescence intensities are plotted as white circles. **(E)** Localization of HIF-1α-IDRs-EGFP and HIF-1α-non-IDR-EGFP in THP-1 cells after 8 h of hypoxia and fixation. Scale bar, 5 μm. **(F)** Representative fluorescence images of EGFP and DAPI showing the patterns of HIF-1α mutants. Scale bar, 5 μm. **(G)** Immunofluorescence staining of HIF-1α in THP-1 cells treated with TAK-981 or control solution under hypoxia. Twenty cells randomly selected from three repeated groups were analyzed. Scale bar, 5 μm. * *P* < 0.05. **(H)** The predicted SUMOylation sites in HIF-1α based on the SUMOplot™ Analysis Program. **(I–J)** SUMO 2/3-mediated SUMOylation and immunofluorescence staining in hypoxia-induced THP-1 cells with the HIF-1α K477F or K391F mutation. Scale bar, 5 μm.

Next, we used subasumstat (TAK-981), a small molecule SUMOylation inhibitor, to determine the effect of SUMOylation on the formation of HIF-1α phase-separated droplets. Immunofluorescence analysis of HIF-1α revealed that TAK-981 significantly reduced the density and fluorescence intensity of hypoxia-induced endogenous HIF-1α phase separation droplets in THP-1 cells (**Figure 5G**). In addition, we predicted and scored the SUMOylation sites of HIF-1α with the SUMOplot™ Analysis Program and focused on the top two sites (amino acids 477 and 391) with scores > 0.9 (**Figure 5H**). We constructed HIF-1α full-length plasmids containing EGFP and the K477F and K391F mutants. The SUMOylation of K391F-mutated HIF-1α decreased under hypoxia compared with that of the full-length protein (**Figure 5I**). Additionally, although the K477F mutation led to a lack of large condensates, we still observed abundant droplets in most cells (**Figure 5J**). However, in the K391F mutant group, only a relatively diffuse pattern of EGFP fluorescence was observed. These results show that hypoxia induces the HIF-1α protein to aggregate in the nucleus and form punctate phase-separation droplets regulated by SUMOylation.

### Targeting the YY1–NUSAP1–HIF-1α nexus in macrophages inhibits PCa tumor growth *in vivo*

To explore the therapeutic effects of targeting the YY1–NUSAP1–HIF-1α nexus in macrophages, we employed Discovery Studio software to dock 1729 small molecules against these three proteins. Among them, tenapanor (T7587, CAS *#*1234423-95-0), a sodium/hydrogen exchanger 3 inhibitor currently used to treat hyperphosphatemia in long-term dialysis patients^37^, emerged with a top 10 docking score across all three proteins. Additionally, we utilized web servers AlphaFold, HDOCK, and Rosetta to generate protein-protein interaction complexes, analyzing amino acid properties and atomic spatial information to pinpoint interaction sites. Comparing the spatial positions of protein interaction sites with those of small molecules revealed a spatial clash between tenapanor’s predicted protein-protein interaction domain and the YY1–NUSAP1–HIF-1α interaction domains (**Figure S3A**). Specially, we also observed that tenapanor spatially clashes to the key binding domains (YY1^200-220^ and NUSAP1^240-250^) we verified within the protein complex, suggesting that tenapanor may competitively inhibit YY1–NUSAP1 interaction, thereby disrupting the YY1–NUSAP1–HIF-1α nexus (**Figures 6A**, and **S3B**). To validate the role of tenapanor in the YY1–NUSAP1–HIF-1α nexus, we conducted an IP analysis in THP-1 cells treated with either tenapanor or DMSO. The results confirmed that tenapanor significantly suppressed the binding of YY1 and HIF-1α to NUSAP1 (**Figures 6B, C**).

**Figure 6.**
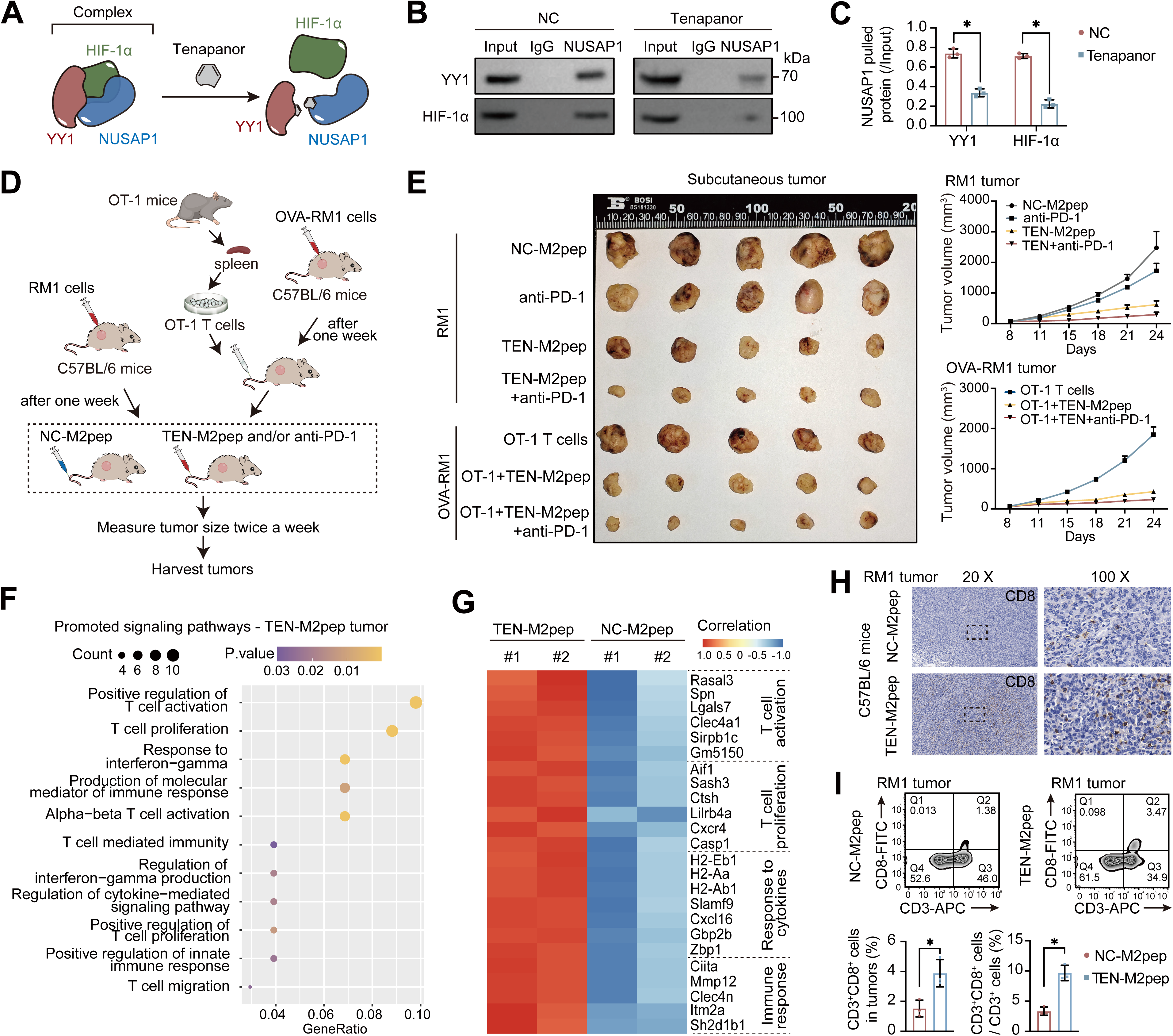
A small molecule inhibitor targeting the YY1–NUSAP1–HIF-1α interaction suppressed PCa progression. **(A)** Schematic diagram showing that tenapanor inhibits the binding of the YY1–NUSAP1–HIF-1α nexus. **(B–C)** Co-IP of NUSAP1 showed that its interactions with YY1 and HIF-1α were suppressed by tenapanor. The protein intensity of YY1 and HIF-1α pulled down from NUSAP1 was analyzed with ImageJ and standardized to that of the input proteins. * *P* < 0.05. **(D–E)** Schematic diagram and tumor growth curve of the effects of TEN-M2pep and control medium on subcutaneous tumorigenesis in mice. **(F–G)** GO analysis based on RNA sequencing data of TEN-M2pep-treated subcutaneous tumors showing enriched immune response-related pathways and representative upregulated genes. **(H–I)** Immunohistochemical staining and flow cytometry of subcutaneous tumors showing the infiltration of CD8^+^ T cells in the indicated groups. * *P* < 0.05.

To verify the effect of tenapanor *in vivo*, we used RM-1 cells to construct subcutaneous tumor-bearing mice (**Figure 6D**). To better target tumor-associated macrophages in the tumor microenvironment, a reported TAM-targeting peptide (M2pep) synthesized on a liposome carrier^38^ was introduced to load tenapanor or the control solvent (TEN-M2pep/NC-M2pep). To clarify its distribution in mouse tissues, we constructed a fluorescent label on TEN-M2pep. Tracer imaging technology further confirmed a noticeable enrichment of TEN-M2pep in mouse subcutaneous tumors within the first two days after injection (**Figure S3C**). Notably, during the one-month observation period after the injection, the growth curve showed that compared with the control treatment, TEN-M2pep significantly hampered tumor growth in the mice (**Figure 6E**). Additionally, we performed RNA sequencing on the harvested tumors, revealing significant enrichment of pathways related to T-cell proliferation and activation in the TEN-M2pep-treated group (**Figures 6F, G**). Meanwhile, we also observed an increased density of tumor-infiltrated CD8^+^ T cells in the TEN-M2pep group by immunohistochemistry and flow cytometry analyses (**Figures 6H, I**).

The efficacy of immune checkpoint inhibitors in tumors largely depends on the quantity of effector T cells in the tumor microenvironment. Herein, we investigated the effect of combination treatment with TEN-M2pep and a PD-1 inhibitor in a PCa subcutaneous xenograft mouse model. The results showed that the PD-1 inhibitor had a limited tumor suppressive effect on the subcutaneous tumor model, whereas the tumors grew significantly more slowly after the combination of the PD-1 inhibitor with TEN-M2pep (**Figure 6E**). Next, we used OVA-RM1 cells expressing ovalbumin (OVA) as a neotumor antigen to construct a mouse subcutaneous PCa tumor model. T cells sorted from OT-1 mouse spleens (OT-1 T cells) were injected into OVA-RM1 tumor-bearing mice to generate tumor-specific exogenous T cells. We found that the combination of OT-1 T cells and TEN-M2pep significantly reduced subcutaneous tumor growth compared to OT-1 T cells alone. Moreover, in the presence of tumor-specific exogenous T cells, a PD-1 inhibitor and TEN-M2pep also showed significant synergistic antitumor effects in this model (**Figure 6E**). Notably, neither TEN-M2pep nor the PD-1 inhibitor caused significant morphological changes in the supportive organs, preliminarily indicating the safety of tenapanor *in vivo* (**Figure S3D**). Flow cytometry analysis of subcutaneous tumors revealed that TEN-M2pep increased the proportion of endogenous CD8^+^ T cells in tumors but had a limited effect on exogenous CD8^+^ T cells from OT-1-treated mice (**Figure S3E**).

### Construction and application of YY1-DbTACs and YY1-DcTACs in an allogeneic mouse transplantation model

Considering the critical role of the YY1–NUSAP1–HIF-1α nexus, we further investigated strategies for impeding the progression of PCa by degrading the YY1 protein. Currently, due to challenges in developing YY1 into a therapeutic agent, there is still a lack of YY1-targeted drugs for clinical use^39^. Proteolytic targeting chimeras (PROTACs) are bifunctional small molecules that can simultaneously bind target proteins and E3 ubiquitin ligases and are a promising strategy for degrading target proteins through the ubiquitin_proteasome system^20^. Hence, utilizing our previously reported covalent DNA framework-based PROTACs (DbTACs) system^40^, we innovatively designed and constructed YY1-targeted DNA-based PROTACs (YY1-DbTACs) to degrade the YY1 protein (**Figure 7A**). This process involved coupling a dibenzo-cyclooctyne (DBCO)-modified DNA strand S1 with an azide-modified E3 ligase ligand through click chemistry. The 5’ end of the DNA strand S2 extended the YY1 binding sequence as the target protein ligand. The above two complementary paired DNA strands self-assembled into YY1-targeted DNA-based PROTACs (YY1-DbTACs) through thermal annealing. The covalent attachment of ligands and the self-assembly of YY1-DbTACs were confirmed by polyacrylamide gel electrophoresis (PAGE) (**Figure S4A**). To investigate the degradation kinetics of YY1-DbTACs *in vitro*, we incubated THP-1 cells with YY1-DbTACs at various concentrations and for various durations. Western blotting confirmed that YY1-DbTACs exhibited concentration- and time-dependent degradation of the YY1 protein (**Figure 7B**). The slight recovery after 36 h may result from counteracting factors, such as the degradation of the DNA double-stranded structure of YY1-DbTACs within 24 h^57^, while YY1 continues to be synthesized. In addition, the degradation mechanism suggested that YY1-DbTACs enhanced YY1 ubiquitination and degradation (**Figure 7C**). As expected, YY1-DbTACs effectively degraded the YY1 protein through the ubiquitination pathway.

**Figure 7.**
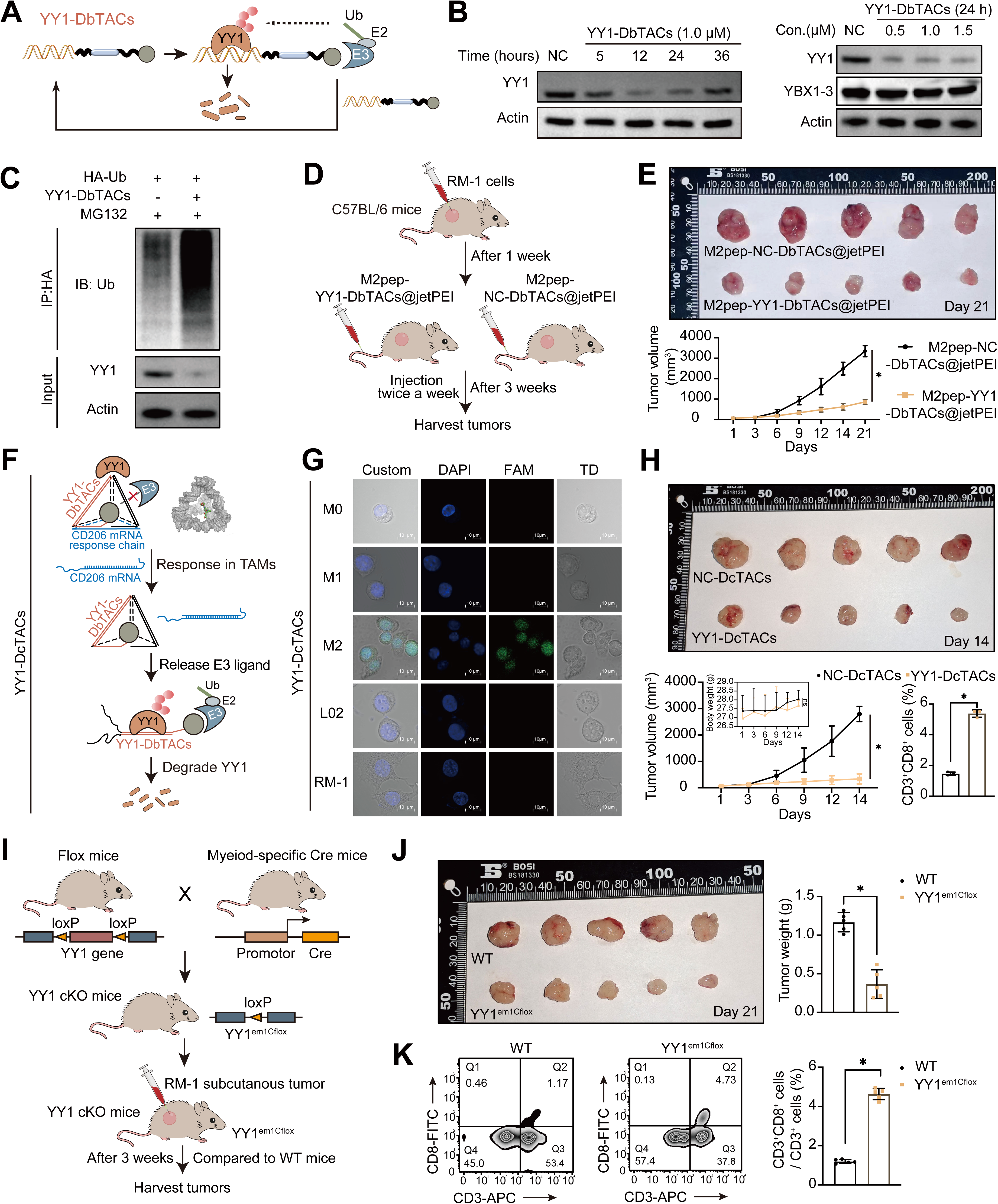
Construction and application of YY1-DbTACs@jetPEI and transgenic mice. **(A)** Schematic diagram illustrating the mechanism of YY1-DbTACs-mediated degradation of the YY1 protein using the PROTAC system. **(B)** Western blot assays showing the amount of YY1 protein in THP-1 cells treated with various concentrations of YY1-DbTACs or incubated for different durations. **(C)** Western blot assays demonstrating that YY1-DbTACs promotes the ubiquitination of YY1 in THP-1 cells. **(D)** Experimental flow chart of the effect of M2pep-YY1/NC-DbTACs@jetPEI on subcutaneous tumorigenesis in mice. **(E)** Tumor volume changes over 21 days following subcutaneous tumor implantation. * *P* < 0.05. **(F)** Schematic diagram illustrating the mechanism of action of YY1-DcTACs based on the PROTAC system and its tetrahedral structure. **(G)** Confocal microscopy images showing the responsiveness of YY1-DcTACs-FAM/BHQ1 in the indicated cell lines. **(H)** Changes in tumor volume within 14 days after injection of YY1/NC-DcTACs via the tail vein in a mouse subcutaneous tumor model. After the subcutaneous tumors were ground into a cell suspension, the T lymphocytes were sorted by flow cytometry, and changes in the ratio of CD3^+^CD8^+^ T cells were observed. * *P* < 0.05. **(I)** Schematic diagram illustrating the construction of YY1 transgenic mice with myeloid conditional knockout and the establishment of a subcutaneous tumorigenesis model. **(J)** Subcutaneous tumor size in mice after 21 days of tumor formation. * *P* < 0.05. **(K)** T lymphocytes were sorted by flow cytometry after grinding the subcutaneous tumors of transgenic mice into a cell suspension, and changes in the ratio of CD3^+^CD8^+^ T cells were observed. * *P* < 0.05.

To investigate the fate of YY1-DbTACs *in vivo*, we injected jetPEI reagent-transfected Cy7-labeled YY1-DbTACs (Cy7-YY1-DbTACs@jetPEI) and free Cy7 dye into the tail veins of allogeneic transplanted mice. Fluorescent images of the mice were collected at different time points after the injection through small animal *in vivo* imaging. The fluorescence intensity at the tumor site of mice injected with Cy7-YY1-DbTACs@jetPEI significantly increased during the experimental period. The tumor site of mice injected with Cy7 peaked at 2 h and then continuously decreased (**Figure S4B**). The fluorescence intensity analysis of isolated mouse organs and tumor tissues revealed that compared with Cy7, Cy7-YY1-DbTACs@jetpei was enriched in tumor tissues after 48 h and was metabolized by the liver and kidney (**Figure S4C**).

The primary reason for the difference in metabolism between the two treatment groups may be the strong lipophilicity of the jetPEI carrier. These results suggest that YY1-DbTACs@jetPEI can accumulate in tumors and exhibit targeting ability. To target YY1 in tumor macrophages, M2pep was combined with YY1-DbTACs@jetPEI. RM-1 cells were used to establish a subcutaneous PCa tumor model in mice, and the distribution and role of M2pep-YY1-DbTACs@jetPEI in subcutaneous tumors were observed via tail vein injection (**Figure 7D**). Fluorescence detection in mice showed that supplementation with M2pep did not significantly inhibit the enrichment of YY1-DbTACs in subcutaneous tumor tissues (**Figure S4D**). After the intermittent tail vein injection of M2pep-YY1/NC-DbTACs@jetPEI for 3 weeks, the growth of subcutaneous tumors in the M2pep-YY1-DbTACs@jetPEI group was significantly suppressed (**Figure 7E**).

Despite the development of M2pep-YY1-DbTACs@jetPEI capable of targeting and degrading YY1, the double-stranded DNA structure of YY1-DbTACs is rapidly degraded within 24 h by nucleases in cells, resulting in a higher dosing frequency, lower applicability, and significant challenges in clinical application due to the difficult and costly preparation of M2pep-loaded liposomes. Therefore, YY1-targeted tetrahedral DNA-caged PROTACs (YY1-DcTACs) were designed and constructed to achieve prolonged degradation of the YY1 protein in M2 macrophages (**Figure 7F)**. This tetrahedral DNA nanostructure (YY1-DcTACs) contains a CD206 mRNA response switch to release YY1-DbTACs in M2 macrophages.

Specifically, an azide-modified E3 ligand is coupled to the DBCO-modified nucleotide site inside the tetrahedron, while the other side of the reported DNA tetrahedral framework is replaced by the binding sequence of YY1 as the target protein ligand. The cage structure of the tetrahedron insulates the E3 ubiquitin ligand to prevent its binding to the E3 ubiquitin ligase, thus blocking YY1-DbTACs. The complementary DNA strand S1 of the modified nucleic acid site was designed as an antisense sequence against CD206 mRNA. Triggered by CD206 mRNA in TAMs, DNA strand S1 dissociates, opening the cage-like structure of the tetrahedron and releasing YY1-DbTACs. The formation of the intact tetrahedron was confirmed by PAGE analysis (**Figure S4E**). To further test the stability of DbTACs under physiological conditions, YY1-DcTACs were incubated in 10% FBS-containing medium for 24 h, and the results showed that YY1-DcTACs did not degrade significantly (**Figure S4F**). To investigate the response efficiency of YY1-DcTACs to CD206, we introduced CD206 mRNA into YY1-DcTACs for strand displacement. The free energy required for the self-assembly (I) and responsive strand displacement (II) of YY1-DcTACs was calculated. The ΔG value between them indicated that YY1-DcTACs could spontaneously respond to CD206 mRNA and undergo strand displacement. Notably, the PAGE results confirmed that S1 and CD206 mRNA were bound at a 1:1 ratio, while the YY1-DcTACs completely dissociated (**Figure S4G**). To further investigate the response efficiency of YY1-DcTACs in living cells, a pair of base pairs on the S1 chain and its complementary chain were modified with FAM fluorophores and BHQ1 quenched groups, respectively, and these DNA strands self-assembled to form modified YY1-DcTACs (YY1-DcTACs-FAM/BHQ1). The fluorescence of YY1-DcTACs-FAM/BHQ1 was quenched due to the close spatial distance between BHQ1 and FAM (< 3 nm). In TAMs, the S1 sequence modified with BHQ1 dissociated upon complementarity with CD206 mRNA, and the fluorescence of YY1-DcTACs-FAM/BHQ1 was restored (**Figure S4H**). The fluorescence pattern confirmed that the fluorescence intensity of YY1-DcTACs-FAM/BHQ1 recovered significantly after the addition of CD206 mRNA (**Figure S4I**). Confocal images revealed that YY1-DcTACs-FAM/BHQ1 responded only to M2 macrophages and not to normal L02 cells, RM-1 cells, M0 macrophages, and M1 macrophages (**Figure 7G**). To further investigate the response efficiency of YY1-DcTACs *in vivo*, the FAM/BHQ1 fluorophore/quencher pairs were substituted with CY5/BHQ3 fluorophore/quencher pairs (ResDcTACs-CY5/BHQ3). ResDcTACs-CY5/BHQ3, YY1-DcTACs labeled with CY5 (DcTACs-CY5), and nonresponsive YY1-DcTACs labeled with the CY5/BHQ2 pair (nonResDcTACs-CY5/BHQ3) were injected into the tail veins of allogeneic transplanted mice. Immunofluorescence colocalization analysis demonstrated that YY1-DcTACs strongly colocalized with CD206 at the tumor site (**Figure S4J**). These results indicate that YY1-DcTACs can specifically induce the release of YY1-DbTACs in tumor tissues through CD206 mRNA. The fluorescence imaging results demonstrated that the fluorescence intensity at the tumor site in the mice injected with DbTACs-CY5 peaked at 3 h and then continuously decreased slowly (**Figure S4K**). Using a similar subcutaneous tumorigenesis model and NC/YY1-DcTACs tail vein injection treatment described above, we observed a remarkable tumor inhibitory effect of YY1-DcTACs until the 14th day (**Figure 7H**). Additionally, the two groups demonstrated no significant difference in body weight. Furthermore, upon grinding, we conducted a flow cytometric examination of subcutaneous tumors in both groups and noted a significant increase in the proportion of CD3^+^CD8^+^ T cells in the YY1-DcTACs group (5.38% ± 4.78% vs 1.46% ± 0.11%, **Figures 7H, S4L**).

To further assess the influence of macrophage YY1 on subcutaneous tumor development, we engineered conditional knockout C57BL/6J-YY1^em1Cflox^ mice with YY1 specifically depleted in macrophages (**Figure 7I**). Through the genetic characterization of bone marrow-derived macrophages (BMDMs) and cells obtained from mouse tails, we clearly delineated the marked suppression of YY1 expression in macrophages compared to that in somatic cells (**Figure S4M**). Subsequent subcutaneous tumor experiments conducted on C57BL/6J-YY1^em1Cflox^ and wild-type mice revealed a noteworthy inhibition of tumor growth in the transgenic mice (**Figure 7J**). Flow cytometry analysis revealed that conditional knockout of YY1 markedly elevated the ratio of CD8^+^ cytotoxic T cells to CD3-labeled T cells within subcutaneous tumors (**Figure 7K**).

In conclusion, we confirmed that suppressing YY1 levels in macrophages effectively restrains PCa growth *in vivo* through the conditional knockout of the YY1 gene or targeted YY1 protein degradation. Notably, tetrahedral nanostructures employing PROTAC technology may pave the way for material advancements toward clinical applications targeting YY1.

## DISCUSSION

YY1, a highly evolutionarily conserved nuclear transcription factor, exerts regulatory control over approximately 7% of human genes^41^. The significance of YY1 extends to various facets of tumor progression, including cell proliferation, DNA repair, epigenetic modification, and tumor immune regulation^42-44^. Our previous study revealed notable expression of YY1 in macrophages and a significant association between increased YY1 expression in TAMs and poorer prognosis in PCa patients^45^. In the present study, we initially observed an elevated ratio of YY1^+^ macrophages within hypoxic regions, and we further elucidated the role of YY1 in promoting the infiltration of macrophages into hypoxic areas and stabilizing the HIF-1α protein. Therapeutic interventions targeting macrophage YY1, along with the use of a transgenic mouse model with conditional knockout of YY1 in macrophages, demonstrated remarkable efficacy in subcutaneous tumor-bearing mouse models. These findings unveil promising avenues for clinical strategies for PCa treatment.

Phase separation is a mechanism that explains the process of multivariate mixtures aggregating into subcellular structures to regionalize synergistically acting molecules, thus accelerating downstream transcription^46^. The occurrence of phase separation typically hinges on the presence of IDRs and protein oligomerization binding regions within the amino acid sequence of a protein^47^. In this study, we also discovered that YY1, NUSAP1, and HIF-1α possess IDR domains, rendering them capable of undergoing phase separation. These three proteins can colocalize and form phase-separated droplets within the cell nucleus, especially under hypoxic conditions. Knocking out the functional fragments within the IDR region markedly diminished the occurrence of phase separation. These findings suggest that the phase separation mechanism plays a crucial role in the YY1-mediated stabilization of HIF-1α. Additionally, the formation and stability of phase separation are regulated by various factors, including protein posttranslational modifications (PTMs)^48^. Herein, we discovered that phosphorylation of YY1 and SUMOylation of HIF-1α enhances their phase separation. Moreover, mutations at relevant protein modification sites reduced the formation of phase-separated droplets, directly impacting protein function. It has been reported that PTMs, such as phosphorylation, ubiquitination, and SUMOylation, can modify the charge properties of amino acid residues, alter local structures, and facilitate intermolecular charge interactions, thereby regulating the intracellular formation of phase separation^49, 50^. Our study highlights the crucial role of epigenetic modification in the mechanism of phase separation in YY1 and HIF-1α.

Hypoxia and overexpression of HIF-1α are involved in tumor immune escape and promote tumorigenesis^7^. The ubiquitin_proteasome system is the classical pathway involved in the stabilization of HIF-1α^51^. Under normoxic conditions, HIF-1α undergoes hydroxylation by prolyl hydroxylases, leading to its recognition by the von Hippel_Lindau E3 ligase (VHL), ubiquitination, and subsequent proteasomal degradation. VHL orchestrates the swift degradation of HIF-1α under normoxia. Conversely, in hypoxic environments, the oxygen-independent slow degradation of HIF-1α further modulates its stability^52^. As a posttranslational modification akin to ubiquitination, SUMOylation affects the ubiquitination process and subsequent substrate degradation^53^. SUMOylation modifications can potentially occupy protein ubiquitination sites, thereby reducing protein ubiquitination levels and stabilizing proteins. Studies have revealed that RSUME^54^ and Cbx4^55^ can enhance HIF-1α SUMOylation, promoting its stability and transcriptional activity under hypoxia. In this study, we report for the first time that NUSAP1 can also modify the HIF-1α protein through its SUMOylation, thus inhibiting HIF-1α ubiquitination and increasing its stability. NUSAP1, a microtubule-associated protein related to mitosis, is implicated in various biological processes, including tumorigenesis^56^. Notably, NUSAP1 contains an SAP domain, which is known to be pivotal for regulating SUMO ligase activity^33^. The SUMOylation sites of HIF-1α are rare, with only a few studies indicating that the C-terminus of HIF-1α, particularly residues K391 and K477, may serve as the primary cluster of SUMOylation sites; previous studies have indicated that mutations at both sites can diminish HIF-1α expression within the nucleus and its transcriptional activity^55, 57, 58^. However, research delineating the disparity between these two sites remains scarce. Interestingly, we discovered that K391 might exert a more significant effect than K477 on inducing HIF-1α SUMOylation-mediated phase separation.

Due to the structural heterogeneity of YY1 and the lack of active sites, few YY1 inhibitors have been applied for clinical tumor treatment^39^. PROTACs are heterodifunctional molecules that utilize the cell’s ubiquitin_proteasome system to degrade targeted proteins of interest, enabling the degradation of "undruggable" proteins ^20, 59^. In this study, we introduced a cage-like tetrahedral nucleic acid structure, YY1-DcTACs, based on the PROTAC technique, which can selectively release DNA-based PROTACs (YY1-DbTACs), effectively degrading the YY1 protein in macrophages and demonstrating its antitumor effects. Furthermore, we have shown that YY1-DcTACs exhibit significant advantages over YY1-DbTACs in terms of structural design and functional characteristics. Specifically, the unique rigid tetrahedral structure of YY1-DcTACs endows them with a longer drug residence time than YY1-DbTACs, thus significantly enhancing therapeutic efficacy. Additionally, the specific response mechanism of YY1-DcTACs, which specifically degrades the YY1 protein in macrophages, provides high precision. These unique properties exhibit tremendous potential in clinical applications, enabling therapeutic effects at lower effective doses while extending treatment intervals and reducing treatment frequency and potential side effects. However, the drug loading capacity of YY1-DcTACs is relatively limited, with each DcTAC molecule effectively releasing only one DbTAC molecule, which limits its therapeutic potential to some extent. Overall, YY1-DcTACs present new prospects for targeting "undruggable" YY1 in clinical therapy for PCa, offering innovative treatment approaches. We anticipate that integrating PROTACs with nanonucleic acids will yield promising solutions for future researchers in clinical tumor treatment.

In conclusion, we observed that YY1^+^ macrophages infiltrate the hypoxic regions of PCa cells. Hypoxia triggers YY1 phosphorylation in macrophages, inducing its phase separation. Concurrently, YY1 interacts with NUSAP1 and promotes the SUMOylation of HIF-1α, inhibiting its ubiquitination and thus stabilizing HIF-1α expression. *In vivo* experiments confirmed that therapy targeting YY1 or the combination of YY1–NUSAP1–HIF-1α inhibits subcutaneous tumor growth in mice. Our study provides insights into the mechanisms of YY1 and HIF-1α in TAMs and provides a solid theoretical basis for the advancement of PCa treatment.

## MATERIALS AND METHODS

### Human tissue samples and tissue microarray

Paraffin-embedded PCa tissue samples were collected from patients who underwent radical prostatectomy between August 2019 and August 2023 in Southeast University Zhongda Hospital. Patients with distant metastasis or receiving androgen deprivation therapy, immune therapy, radiotherapy, or chemotherapy were excluded. At least two pathologists confirmed the pathological diagnosis. A tissue microarray was constructed using 47 tissue cores (Prostate cancer tissue) obtained from tumor centers of different PCa tissues and 30 normal gland tissue cores (Para-cancerous tissue) adjacent to cancerous regions. Each core had a diameter of 1 mm. One core from the Prostate cancer tissue and four cores from the Para-cancerous tissue were excluded due to the tissue area being less than 50% of the designated area. The baseline information of included patients was listed in **Table S1**. The collection and use of clinical samples are completed under the supervision of Ethics Committee of Zhongda Hospital Southeast University (approval ID 2022ZDKYSB099).

### Imaging mass cytometry (IMC)

After slicing the tissue microarray continuously, one slice was chosen for hematoxylin and eosin (H&E) staining. The histological type of each core was verified by two pathologists. Following this, the labeled sections were stained with IMC target antibodies containing metal labels. These pre-sections were then scanned using the Hyperion Imaging System (Fluidigm) to generate multiplexed images. Subsequently, the acquired data underwent analysis through several steps: spillover signal compensation, image denoising, image contrast enhancement, and cell segmentation. The methodology described above is based on previously published research^60^. Furthermore, to identify the local accumulation of a specific cell type and its neighboring cells, the patch and milieu detection algorithm^61^ was employed to investigate the correlation among YY1, CD68, and HIF-1α.

### Cell lines and culture

Human monocyte THP-1 and mouse macrophage RAW264.7 cell lines were provided by the American Type Culture Collection (ATCC, USA). Cells were incubated in the RPMI 1640 medium or DMEM medium (Gibco, Thermo Fisher Scientific, Waltham, Massachusetts, USA) containing 10% fetal bovine serum (FBS, Gibco), 1% penicillin G and streptomycin sodium (Gibco). For normal conditions, cells were cultured at 37°C with 5% CO_2_ in a humidified incubator. For hypoxic conditions, cells were cultured at 37°C with, 5% CO_2_, adding cobalt chloride (CoCl_2_) into the medium.

### Reagents, antibodies and plasmids

Reagents and antibodies used in this study include: YY1 (Abcam, cat. ab109237, 1:1000 for western blot (WB), 1:200 for immunofluorescent (IF) staining); YY1 (H-10) (Santa Cruz, cat. sc-7341, 1:50 for immunoprecipitation (IP), 1:200 for IF); NUSAP1 (Proteintech, cat. 12024-1-AP, 1:2000 for WB, 1:100 for IP, 1:200 for IF); HIF-1α (abcam, cat. ab51608, 1:1000 for WB, 1:500 for IF, 1:100 for IP); Lyn (Proteintech, cat. 18135-1-AP, 1:1000 for WB); and Tubulin (Proteintech, cat. 11224-1-ap, 1/2000 for WB); SUMO 2/3 (Proteintech, cat. 11251-1-ap, 1:1000 for WB); Ubiquitin (Proteintech, cat. 10201-2-ap, 1:1000 for WB); Flag (Cell signaling Technology, cat. 14793S, 1:1000 for WB, 1:200 for IP); EGFP (Proteintech, cat. 66002-1-lg, 1:2000 for WB, 1:200 for IP); HA (Proteintech, cat. 51064-2-ap, 1:2000 for WB, 1:200 for IP); Phospho-Serine (Immunechem, cat. ICP9806, 1:250 for WB); Phospho-Tyrosine (Immunechem, cat. ICP9805, 1:250 for WB); Phospho-YY1 (Affinity, cat. AF3694, 1:1000 for WB); GAPDH (Acton, cat. 10R-G109A, 1:1000 for WB); F4/80 (Aifang biological, cat. SAF002, 1:500 for mIHC); Goat anti-mouse IgG-HRP (Santa Cruz, cat. sc2005,1:5000 for WB); Goat anti-rabbit IgG-HRP (Santa Cruz, cat. sc-2004, 1:5000 for WB); PE-CD8 (BD Bioscience, cat. 553030, 1:50 for flow cytometry (FC)); APC-CD3 (BD Bioscience, cat. 555275, 1:50 for FC); PE-CD274 (Biolegend, cat. 329706, 1:50 for FC); 1,6-hexanediol (Aladdin, cat. H103708); Tenapanor (TargetMol, T7587, cat. 1234423-95-0); MG132 (MACKLIN, cat. HY132-59); CHX (MACKLIN, cat. HY-12320, 100 µg/ml for CHX assay); Subasumstat (MACKLIN, TAK-981, cat. HY-111789); Na_3_VO_4_ (MACKLIN, cat. 13721-39-6); SU6656 (MACKLIN, cat. HY-B078); K3PO4 (MACKLIN, cat. NONE8222); Protein A/G Magnetic Beads (Beyotime, cat. P2108); Anlotinib (cat. AL3818) dihydrochloride (Aladdin, cat. 1360460-82-7); Cobalt Chloride Solution (MACKLIN, cat. 7791-13-1); anti-mouse PD-1 (Bio X Cell, cat. 857122D1); Anti-mouse lgG (Bio X Cell, cat. 84932331); The full-length coding sequences of YY1, HIF-1α and NUSAP1 (or their variants), as well as IDRs of YY1 and HIF-1α were individually subcloned into a modiffed version of a pGEX vector with 6 × His and EGFP or mCherry at the N-terminus. A bacterial expression system was used to express these coding sequences and recombinant proteins were purified using Ni-NTA agarose. Meanwhile, the full lengths of YY1, HIF-1α and their mutants, as well as NUSAP1 were individually subcloned into a eukaryotic EGFP or mCherry expression vector for studying phase separation. The full-length and truncated coding sequences of YY1, HIF-1α, and NUSAP1 were also individually subcloned into eukaryotic expression vectors N-terminal 3×HA-tag, 3×His-tag and 3×Flag-tag at the N-terminus for studying protein binding.

### Gene knockout system

Small interfering RNAs (siRNAs) or short hairpin RNAs (shRNAs) targeting YY1, HIF-1α, and NUSAP1 were purchased from GenePharma Co. (Shanghai, China). The sequences are listed in **Table S2**. CRISPR/Cas9 YY1 knockout system was constructed by GeneChem.

### RNA extraction and RT-PCR

An RNA extraction kit (Takara Kusatsu) was used for RNA extraction according to the manufacturer’s protocol. Complementary DNA (cDNA) was prepared by reverse transcription polymerase chain reaction (RT-PCR) from purified RNA with the Hiscript II First-Strand cDNA Synthesis Kit (Vazyme). Quantitative Real-time PCR was performed using the MonAmpTM SYBR Green qPCR Mix (Monad Biotech). The comparative threshold cycle method was used to calculate the relative gene expression levels by normalizing to the reference gene GAPDH.

### Western blotting

Total protein was extracted from cells by using the RIPA lysis buffer (KeyGene Biotech, 1:1000) according to the manufacturer’s instructions. The protein concentration was determined with the bicinchoninic acid assay (BCA, KeyGene Biotech). The protein was separated by sodium dodecyl sulfate-polyacrylamide (SDS) gel electrophoresis and transferred onto a polyvinylidene difluoride membrane (PVDF, Merck Millipore). The membrane was then blocked for 1 h with 5% skimmed milk and incubated with the following primary antibodies overnight followed with horseradish peroxidase (HRP) conjugated secondary antibodies.

### RNA-FISH assay combined with immunofluorescence

Cells were treated by the procedure of immunofluorescence experiment as described above, and then subjected to the following RNA FISH assay. A set of Stellaris FISH probes was designed using the Stellaris™ Probe Designer software (Biosearch Technologies), and the Cy3-labeled probes were synthesized by Servicebio. Cell slides were fixed in in situ hybridization fixative for 20 min, and washed thrice with PBS (PH7.4), and washed once with 70% ethanol. Prior to hybridization, the pre-hybridization solution was added dropwise and incubated at 37 ° C for 1 h. Then dump the pre-hybridization solution, drop the probe-containing hybridization solution, and mix overnight in the incubator. Cells were then washed once in the washing buffer, 2 × SSC, 37 ° C for 10min, 1 × SSC, 37 ° C for 2 × 5min, 0.5 × SSC 37 _ for 10 min. Finally, the nuclei were stained with DAPI and imaged using the FV3000.

### Immunoprecipitation-mass spectrometry (IP-MS) and coimmunoprecipitation assays (CO-IP)

For IP-MS assays, the Protein A/G was pre-incubated with the indicated YY1 or rabbit IgG (Proteintech, cat. B900610) for 4 hours. The cell lysis supernatant was then incubated with antibody-coupled magnetic beads at 4_ overnight. Elute with 0.1M Glycine-HCl, pH3.0 and neutralize with 0.5M Tris-HCl, pH7.4, 1.5M NaCl for protein mass spectrometry analysis (GeneChem). For Co-IP assays, the extracts were immunoprecipitated with anti-Flag or anti-HA magnetic beads (Thermo Fisher Scientific) overnight at 4_ and boiled with SDS loading buffer. The released binding proteins were then eluted for western blotting analysis using the indicated antibodies.

### Ubiquitination assay

Cells were incubated with proteasome inhibitor MG132 (50_μg/ml) for 6_h in 37_°C incubator and then lysed with IP lysis buffer. Lysates were then incubated with HIF-1α antibody (2_μg, Abcam) as well as IgG (rabbit) coupled to protein A/G magnetic beads at 4 °C overnight to pull down the HIF-1α proteins. After washing with PBS for five times, proteins were denatured at 100_°C for 5_min and then separated by 10% SDS-PAGE. Ubiquitin antibody was used to detect the ubiquitination of HIF-1α.

### SUMOylation assay

Cells were washed with cold wash buffer (10 mM NEM in PBS) and then lysed with lysis buffer (protease inhibitor and phosphatase inhibitor cocktails). Cell lysates were sonicated until they became fluid and then diluted 1:10 with dilution buffer (protease inhibitor and phosphatase inhibitor cocktails). Lysates were then incubated with anti-HIF-1α antibody coupled to protein A/G magnetic beads at 4 °C overnight. Beads were boiled in SDS sample buffer. SUMOylated HIF-1α was detected by western blot using an anti-SUMO2/3 antibody.

### Immunofluorescence staining

Cells were seeded on glass coverslips in glass bottom dish (φ14 mm) and cultured overnight. Subsequently, cells were fixed with the Immunol Staining Fix Solution (Beyotime) for 30 min at room temperature. After blocking by 10% FBS for 30 min at room temperature, cells were incubated with a primary antibody for 30 min at room temperature. After washing thrice with PBS, cells were incubated with an Alex-Fluor-488 or 594-conjugated secondary antibody (Beyotime) for 30 min at room temperature. Finally, after washing thrice with PBS, nuclei were counterstained with DAPI (Beyotime), and images were captured by Fluoview FV3000 fluorescence confocal microscope.

### Cell imaging, phase separation and fluorescence recovery after photobleaching (FRAP) assays

Cells were cultured in a glass bottom dish. Before Cell imaging, the culture medium was exchanged for Opti-MEM. Images were acquired by Fluoview FV3000 fluorescence confocal microscope with 60× oil immersion lens. Fluorescence signals were obtained with excitation/emission at 488/546 nm for EGFP. The droplet fusion was recorded for 20 to 30 time points (200 to 300 s). The intrinsically disordered regions (IDRs) of YY1, NUSAP1 and HIF-1α proteins were predicted by the PONDR (Predictor of Natural Disordered Regions) website, and the VSL2 algorithm was used in the prediction to obtain the disordered maps of the three proteins. The HIF-1α-intrinsically disordered regions (IDR) 1-EGFP (amino acids 1-80), HIF-1α-IDR2-EGFP (amino acids 391-750), HIF-1α-non-IDR-EGFP (amino acids 81-390), HIF-1α-whole-EGFP (amino acids 1-826), NUSAP1-whole-mCherry (amino acids 1-441) and YY1-whole-mCherry (amino acids 1-414) fusion protein was synthesized by GenScript (Piscataway). Potassium phosphate buffer (pH 7.0) was used for desalinization of the fusion proteins. Polyethylene glycol (PEG) 8000 (30%), as a molecule crowder, together with desalted fusion protein form solutions with different protein concentration gradients (1, 2, 5, 10, and 20 μM). The mixed solution was loaded onto glass slides with coverslips, followed by immediate visualization by a Fluoview FV1000 fluorescence confocal mif croscope (Olympus).

The fluorescence recovery after photobleaching (FRAP) assay was performed on Fluoview FV3000 fluorescence confocal microscope with 60× oil immersion lens. The transfected 293T cells were cultured in a glass bottom dish. Before FRAP, the culture medium was exchanged for Opti-MEM. The droplets in cells were bleached for 3 to 5 s with 10 to 20% of the maximum laser power of a 488 nm laser (1 AU). The recovery was recorded for 20 to 30 time points after bleaching (200 to 300 s). The fluorescent intensity of the bleached area over time was calculated by Zen. During image acquisition, cells were incubated in a chamber at 37◦C supplied with 5% CO_2_.

### Mice tumorigenicity assay

C57BL/6 male mice were provided by the Comparative Medical Center of Yangzhou University and raised at the Animal Center of Southeast University under the standard conditions. Before the experiment, the mice adapted to the laboratory environment for 2-3 weeks until 6-8 weeks old. All operations on experimental mice are performed under the supervision of the Animal Ethics Committee of Southeast University. RM-1 cells (5×10^6^) alone or ones mixed with RAW264.7 cells (5×10^6^) transfected with si/nc-YY1, were injected subcutaneously on the back of mice. M2pep (peptide YEQDPWGVKWWY)^38^, which was synthesized by RuixiBio (Xi’an, China), was modified on a liposome carrier and loaded with Tenapanor or control solvent to construct Ten-M2pep or NC-M2pep. In the in vivo therapeutic experiments, Ten-M2pep or NC-M2pep were injected intravenously one week after the subcutaneous tumor construction. Tumor volume was evaluated twice a week after injection following the formula of length×width^2^/2.

### Flow cytometry

Fresh tissue was cut into pieces and cultured in Collagenase IV and DNase I for 30 min at room temperature. To quantify the cell subgroup and the expression of molecules on the cells, the purified single-cell suspensions were blocked with mouse FcR blocking reagent (Miltenyi Biotec) for 10_min at 4_°C prior to surface staining. Purified cells were stained with PE-CD8 and APC-CD3 at 4 °C for 30 min. Dead cells were excluded with eFluor506. The gate of positive cells for each dye were set based on single-stained cells and isotype controls. Flow cytometry analysis was performed using a FACS flow cytometer (BD Biosciences) and analyzed by FlowJo V10 (TreeStar).

### Fabrication of YY1-DbTACs

All DNA strands were purchased from Biolink Biotechnology. Single-stranded DNA S1 (1000 μL, 50 μM) was mixed with CRBN ligand solution (50 μL, 1000 μM). The solution was shaken for 2 hours (37°C, 400 rpm). Subsequently, the mixture was combined with pre-designed single-stranded DNA S2 (1:1 molar ratio). The resulting solution was heated for 5 min (95°C) using a T-series multi-block thermal cycler (LongGene), then annealed for 30 min (4°C), resulting in the YY1-DbTACs structure. Finally, the prepared YY1-DbTACs were stored at 4°C.

### Fabrication, self-assembly and stability of YY1-tetrahedral-DNA-nanostructure (YY1-TDN)

For YY1-TDN, single-stranded DNA S2 (1000 μL, 50 μM) was mixed with a solution of CRBN ligand (50 μL, 1000 μM). The solution was shaken for 2 hours (37°C, 400 rpm). Subsequently, the mixture was combined with the pre-designed single-stranded DNA sequences S1, S3, and S4 in a 1:1:1:1 molar ratio. The resulting solution was heated for 10 min (95°C) using a PCR machine (LongGene) and then annealed for 35 min (4°C) to obtain the YY1-TDN structure. Finally, the prepared YY1-TDN was stored at 4°C.

To verify the covalent binding of ligands and the principle of stepwise self-assembly of YY1-TDN, a PAGE (10%) gel was employed. 5 μL of each sample were mixed with 1 μL of DNA loading buffer (6×). Electrophoresis was conducted in 1× TBE-Mg2^+^ buffer at 120V for 70 min. The gel was stained with SYBR Green (10,000×, Solarbio Science & Technology) in the dark for 0.5 hours, and then the images were captured using a Bio-RAD Gel Doc EZ Imager (Hercules). YY1-TDN (2 μM) was mixed with PBS and IMDM medium containing 10% FBS, and incubated at 37°C. Samples prepared as described above were collected at various time points (1, 2, 4, 6, 12, 24 h), and analyzed for the stability of YY1-TDN using PAGE (10%) gel electrophoresis.

### Response efficiency of YY1-TDN

The YY1-TDN solution was mixed with the complementary RNA strand S1 in equal molar ratios and incubated at 37°C for 6 hours. The samples were then collected and analyzed using 10% PAGE gel electrophoresis and fluorescence spectrophotometer detection. Macrophages of types M0, M1, and M2, normal hepatocytes L02, and cancer cells were cultured in confocal dishes at a density of 1×10^5^ cells per dish. 200 μL of 1 μM YY1-TDN was added to each dish, and the cells were incubated at 37°C under 5% CO_2_ for 6 hours. The culture medium was discarded, and the cells were washed three times with cold PBS buffer for 30 sec. The cells were then fixed with 4% paraformaldehyde at room temperature, followed by three washes with cold PBS buffer for 5 min. The cell nuclei were stained with DAPI (purchased from Jiangsu KeyGEN BioTECH, cat. KGA215-50, 1 μg/mL, 100 μL) for 10 min at room temperature and washed. Images were acquired using a confocal laser scanning microscope (FV1000, Olympus).

### Construction of transgenic mice

In-Fusion cloning method was used to construct a homologous recombinant donor vector, which was verified by enzyme digestion. C57BL/6J-YY1^em1Cflox^ mice was constructed under the following steps. Cas9 mRNA, gRNA and donor vector were injected into the fertilized eggs of C57BL/6J mice by microinjection operating system to obtain F0 generation mice. F1 hybrid mice (YY1^flox/+^) were then obtained by mating F0 mice with C57BL/6J mice. Breeding of YY1^flox/+^ mice is carried out through both self-crossing and hybridization with Cre mice, resulting in obtaining flox homozygous mice (YY1^flox/flox^) and flox heterozygous Lyz2-Cre double positive mice (YY1^flox/+: Cre+^). Homozygous mice were mated with double positive mice, and finally the target mice with flox homozygous Lyz2-Cre positive genotype (YY1^flox/flox: Cre+^) were obtained.

### Statistical analysis

All experiments were repeated three times independently, and Student’s t-test or one-way ANOVA was used to compare the results in the research groups. All statistical analysis was carried out by SPSS software version 16.0, and p<0.05 indicated that there was a statistically significant difference.

## List of Supplementary Materials

Figure S1 to S4 for multiple supplementary figures

Table S1 for supplementary table 1

Table S2 for supplementary table 2 (Excel file)

## Acknowledgments

We are grateful to the staff of the Public Scientific Research Platform of Zhongda Hospital Southeast University, and the Cultivation and Construction Site of the State Key Laboratory of Intelligent Imaging and Interventional Medicine (Southeast University), for technical assistance. The doctoral students in this project were under the support of Southeast University Innovation Capability Enhancement Plan for Doctoral Students.

## Funding

National Natural Science Foundation of China 82203685 (WL)

National Natural Science Foundation of China 92359304 (SJ)

National Natural Science Foundation of China 82330060 (SJ)

National Natural Science Foundation of China 823B2067 (SC)

Fundamental Research Funds for the Central Universities 2242023K5007 (BX)

General social development project of Jiangsu Science and Technology Department BE2023771 (BX)

China Postdoctoral Science Foundation funded project 2023M730572 (WL)

Outstanding Youth Foundation of Jiangsu Province of China BK20220150 (YM)

Open Research Fund of State Key Laboratory of Digital Medical Engineering 2023-K10 (BX)

Top Talent Support Program for young and middle-aged people of Wuxi Health Committee HB2023117 (MZ)

## Author contributions

Conceptualization: BX, SJ, MC

Methodology: WL, SC, YM, RN, CC, CS, YH, XY

Investigation: JL, WM, SZ, MZ, ZC, RN Visualization: WL, TW, YC, YH, NS

Clinical samples collection: SZ, KL, LHZ, LZ

Funding acquisition: BX, SJ, YM, WL, SC, MZ

Project administration: BX, MC

Supervision: BX, SJ

Writing – original draft: WL, YM, SC

Writing – review & editing: BX, SJ, JL, WM, SZ, MZ

## Data and materials availability

All data are available in the main text or the supplementary materials.

## Ethics approval and consent to participate

All human tumor tissue samples were collected in accordance with the national and institutional ethical guidelines. The study design was approved by the Ethics Committee of Zhongda Hospital Southeast University. The approval ID is 2022ZDKYSB099. All animal experimental protocols were approved by the Ethics Committee of Zhongda Hospital Southeast University and conducted following the National Guidelines for the Health Use of Laboratory Animals. The approval ID is 20220225034.

## Consent for publication

All authors approved the manuscript for submission and gave their consent for publication.

