## Supplementary Figures 1-4 and Tables 1-2 for "YY1 Enhances the Stability of HIF-1α Protein by Interacting with NUSAP1 in Macrophages within the Prostate Cancer Microenvironment": Supplementary Figure 2.pdf

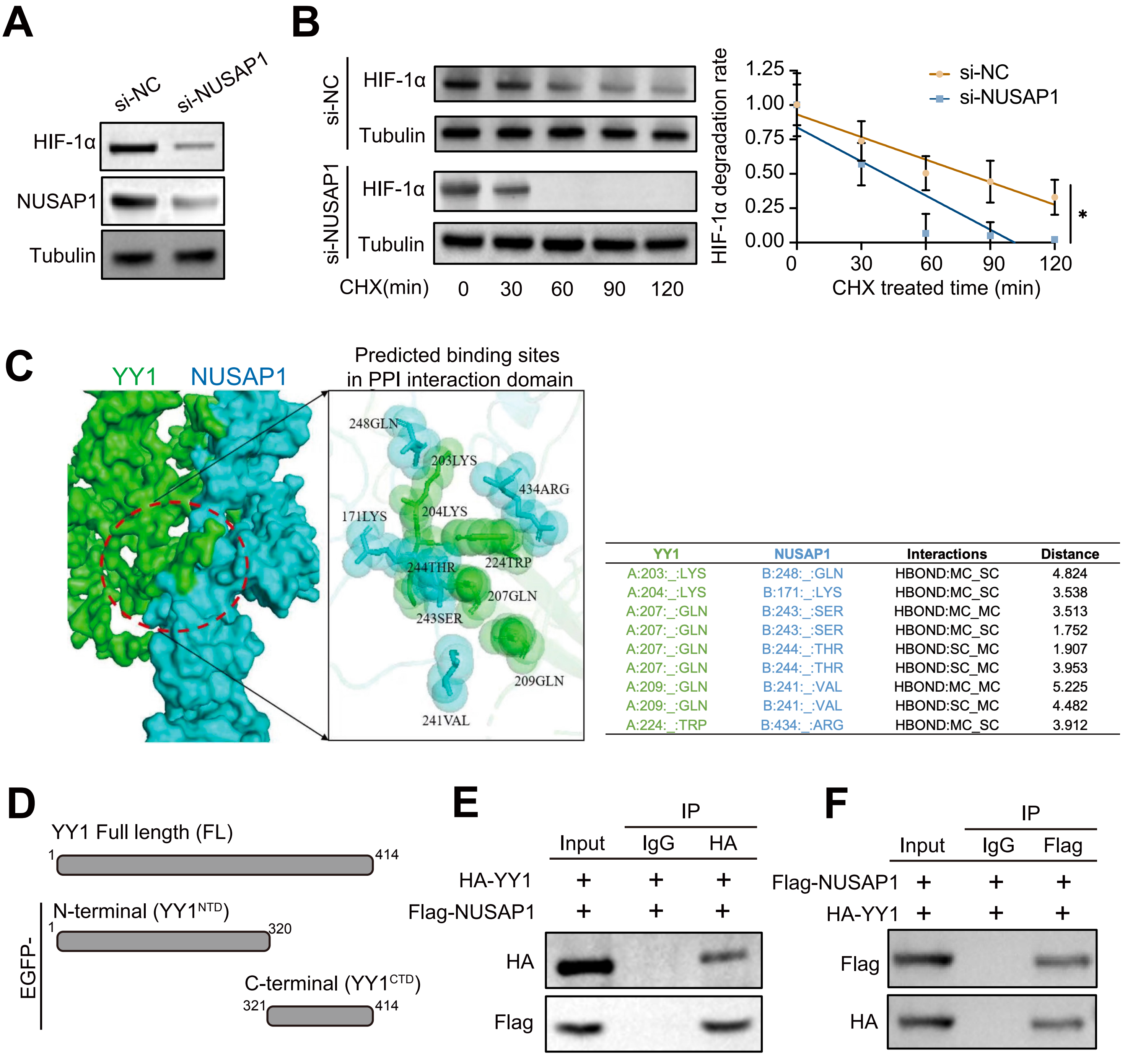


| YY1 | NUSAP1 | Interactions | Distance |
|---|---|---|---|
| A:203:_:LYS | B:248:_:GLN | HBOND:MC_SC | 4.824 |
| A:204:_:LYS | B:171:_:LYS | HBOND:MC_SC | 3.538 |
| A:207:_:GLN | B:243:_:SER | HBOND:MC_MC | 3.513 |
| A:207:_:GLN | B:243:_:SER | HBOND:MC_SC | 1.752 |
| A:207:_:GLN | B:244:_:THR | HBOND:SC_MC | 1.907 |
| A:207:_:GLN | B:244:_:THR | HBOND:SC_MC | 3.953 |
| A:209:_:GLN | B:241:_:VAL | HBOND:MC_MC | 5.225 |
| A:209:_:GLN | B:241:_:VAL | HBOND:SC_MC | 4.482 |
| A:224:_:TRP | B:434:_:ARG | HBOND:MC_SC | 3.912 |
