## Supplementary Figures 1-4 and Tables 1-2 for "YY1 Enhances the Stability of HIF-1α Protein by Interacting with NUSAP1 in Macrophages within the Prostate Cancer Microenvironment": Table S1.pdf

| Table S1 | | |
|---|---|---|
| Characteristics | PCa(N=46) | Para-carcinoma (N=26) |
| Age, year | | |
| Mean±SD | 72.76±8.34 | 70.85±7.34 |
| Median[min-max] | 72.50[56.00,86.00] | 70.50[57.00,86.00] |
| tPSA, ng/mL | | |
| Mean±SD | 29.59±66.95 | 8.37±5.88 |
| Median[min-max] | 11.65[0.72,335.00] | 6.56[0.72,23.30] |
| Gleason Score | | |
| 6 | 14(19.44%) | 26(36.11%) |
| 7 | 14(19.44%) | 0(0%) |
| 8 | 4(5.56%) | 0(0%) |
| 9 | 14(19.44%) | 0(0%) |
| T stage | | |
| 1 | 1(1.39%) | 3(4.17%) |
| 2 | 25(34.72%) | 23(31.94%) |
| 3 | 20(27.78%) | 0(0%) |
| N stage | | |
| 0 | 38(52.78%) | 26(36.11%) |
| 1 | 8(11.11%) | 0(0%) |
| M stage | | |
| 0 | 46(63.89%) | 26(36.11%) |
| Hypertension | | |
| 0 | 22(30.56%) | 14(19.44%) |
| 1 | 24(33.33%) | 12(16.67%) |
| Diabetes | | |
| 0 | 34(47.22%) | 21(29.17%) |
| 1 | 12(16.67%) | 5(6.94%) |
| Coronary artery disease | | |
| 0 | 38(52.78%) | 23(31.94%) |
| 1 | 8(11.11%) | 3(4.17%) |
| Cerebral infarction | | |
| 0 | 36(50.00%) | 21(29.17%) |
| 1 | 10(13.89%) | 5(6.94%) |
| Smoking | | |
| 0 | 33(45.83%) | 20(27.78%) |
| 1 | 13(18.06%) | 6(8.33%) |
| Drinking | | |
| 0 | 36(50.00%) | 23(31.94%) |
| 1 | 10(13.89%) | 3(4.17%) |

**Table S1. The baseline information of patients included in the IMC analysis.**
