## Supplementary Figures 1-4 and Tables 1-2 for "YY1 Enhances the Stability of HIF-1α Protein by Interacting with NUSAP1 in Macrophages within the Prostate Cancer Microenvironment": Table S2.pdf

**siRNA**

| | |
|---|---|
| si-HIF-1a | GGGAUUAACUCAGUUUGAATT |
| | UUCAAACUGAGUUAAUCCCTT |
| si -LYN | S-GGUGCUAAGUUCCCUAUUATT |
| | AS-UAAUAGGGAACUUAGCACCTT |
| si -NUSAP1 | CUGCUACUAAAGAUAAUGATT |
| | UCAUUAUCUUUAGUAGCAGTT |
| si-YY1 | CUGAGGUAACUCUUCUUGCTT |
| | GCAAGAAGAGUUACCUCAGTT |

**Primer**

| | |
|---|---|
| YY1 | YY1F：CCATCTCAGCCTCCCAAGTAG |
| | YY1-R：TGTACTATGCTTCTCCCACAGGA |
| LYN | LYN_3F：AGGGAGGAGCCCATTTAC |
| | LYN_3R：GCACTTTGCCACCTTCA |
| HIF-1a | HIF1A_F：CTGAGGGGACAGGAGGA |
| | HIF1A_R：CACACGCGGAGAAGAGA |

**Vector**

| Gene | Vector |
|---|---|
| NUSAP1_pcDNA3.1(+) | pcDNA3.1(+) |
| pcDNA3.1-flag-NUSAP1-1-441 | pcDNA3.1(+) |
| pcDNA3.1-flag-NUSAP1-1-158 | pcDNA3.1(+) |
| pcDNA3.1-flag-NUSAP1-159-441 | pcDNA3.1(+) |
| NUSAP1(NM_016359(del240-250aa)) | CV702 |
| YY1(NM_003403-HA) | GV712 |
| YY1(NM_003403(del200-226aa)-HA) | GV712 |
| YY1 1-320_pcDNA3.1(+) | pcDNA3.1(+) |
| YY1(NM_003403(1-294aa)) | GV230 |
| YY1(NM_003403(Y145/185/251/254F)) | GV739 |
| YY1(NM_003403(Y8/383F)) | GV739 |
| YY1(321-414aa)-EGFP | pcDNA3.1(+) |
| YY1(1-320aa)-EGFP | pcDNA3.1(+) |
| YY1(1-414aa)-EGFP | pcDNA3.1(+) |
| PGMLV-CMV-H_HIF1A(amino acid,1to390)-12×His-EF1-ZsGreen1-T2A-Puro | GM-8201 |
| HIF1A(NM_001530(391-826aa)-6His) | GV658 |
| HIF-1a-K477F | pcDNA3.1-EGFP |
| HIF-1a-K391F | pcDNA3.1-EGFP |
| HIF-1a | pcDNA3.1-EGFP |
| HIF-1a △380-752 | pcDNA3.1-EGFP |
| HIF-1a △1-81 | pcDNA3.1-EGFP |
| HIF1a 1-80_pcDNA3.1(+) | pcDNA3.1(+) |
| HIF1a 391-750_pcDNA3.1(+) | pcDNA3.1(+) |
| GM-51562: PCDNA3.1-HA-Ubiquitin(WT) | PCDNA3.1 |
| PGMLV-CMV-H_SUMO3-3×Flag-PGK-Puro | PGMLV-CMV-MCS-3×Flag-PGK-Puro |

**Table S2. siRNA, primers, and vectors.**
