## Supplementary figures and images for "YY1 Enhances the Stability of HIF-1α Protein by Interacting with NUSAP1 in Macrophages within the Prostate Cancer Microenvironment"

### Supplementary Figure 1.pdf

#### Supplementary Figure 1

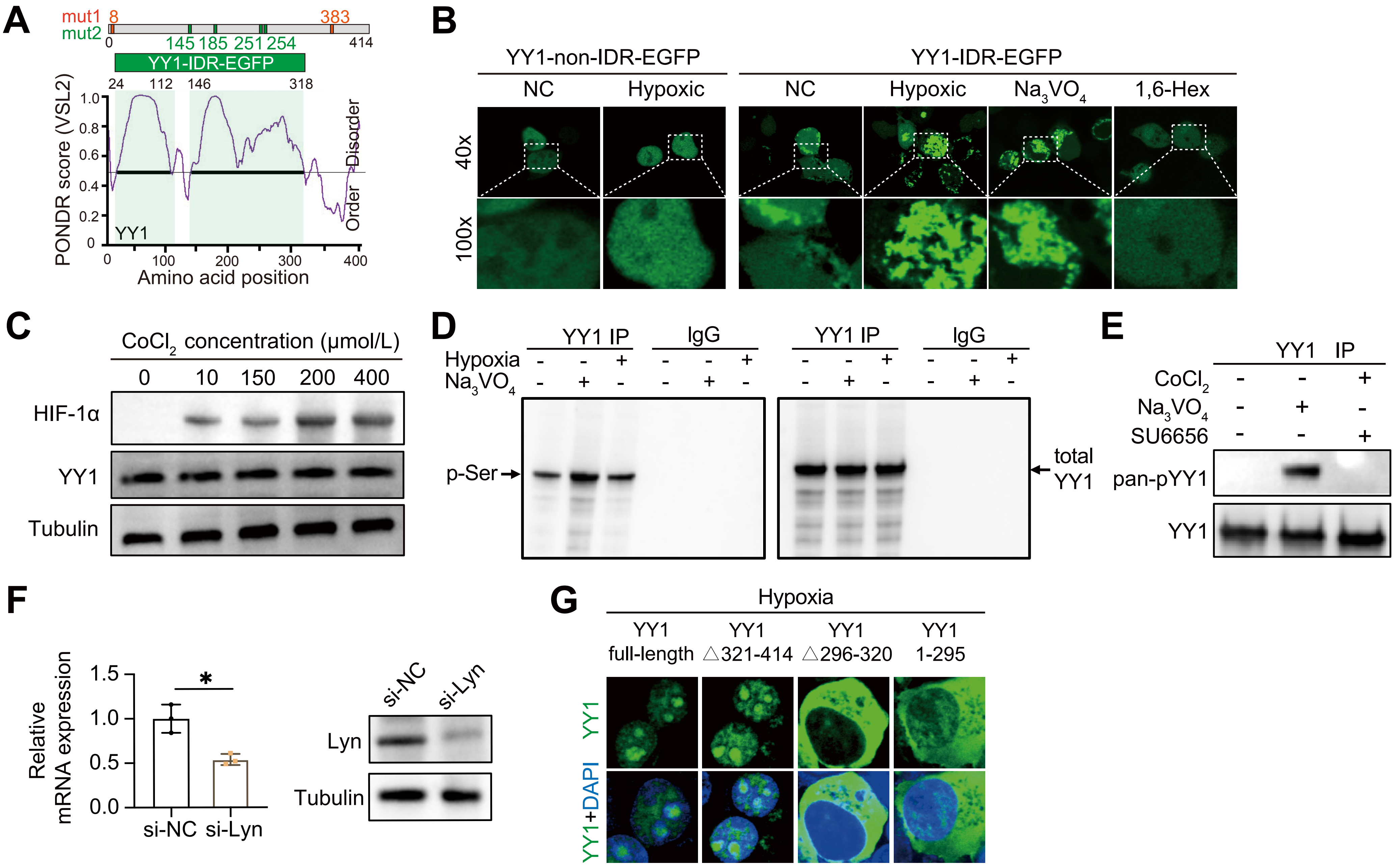

### Supplementary Figure 3.pdf

#### Supplementary Figure 3

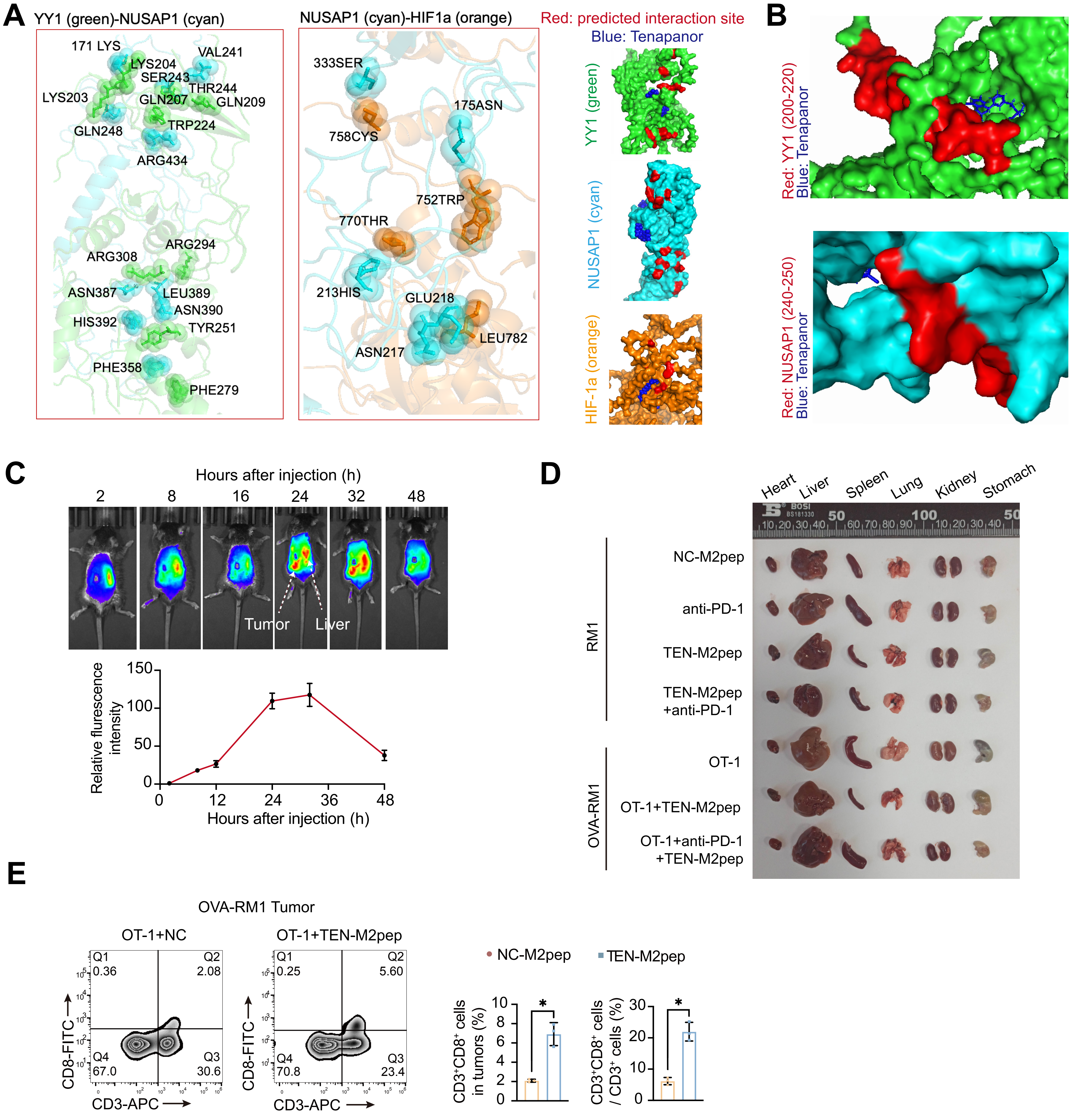

### Supplementary Figure 4.pdf

#### Supplementary Figure 4

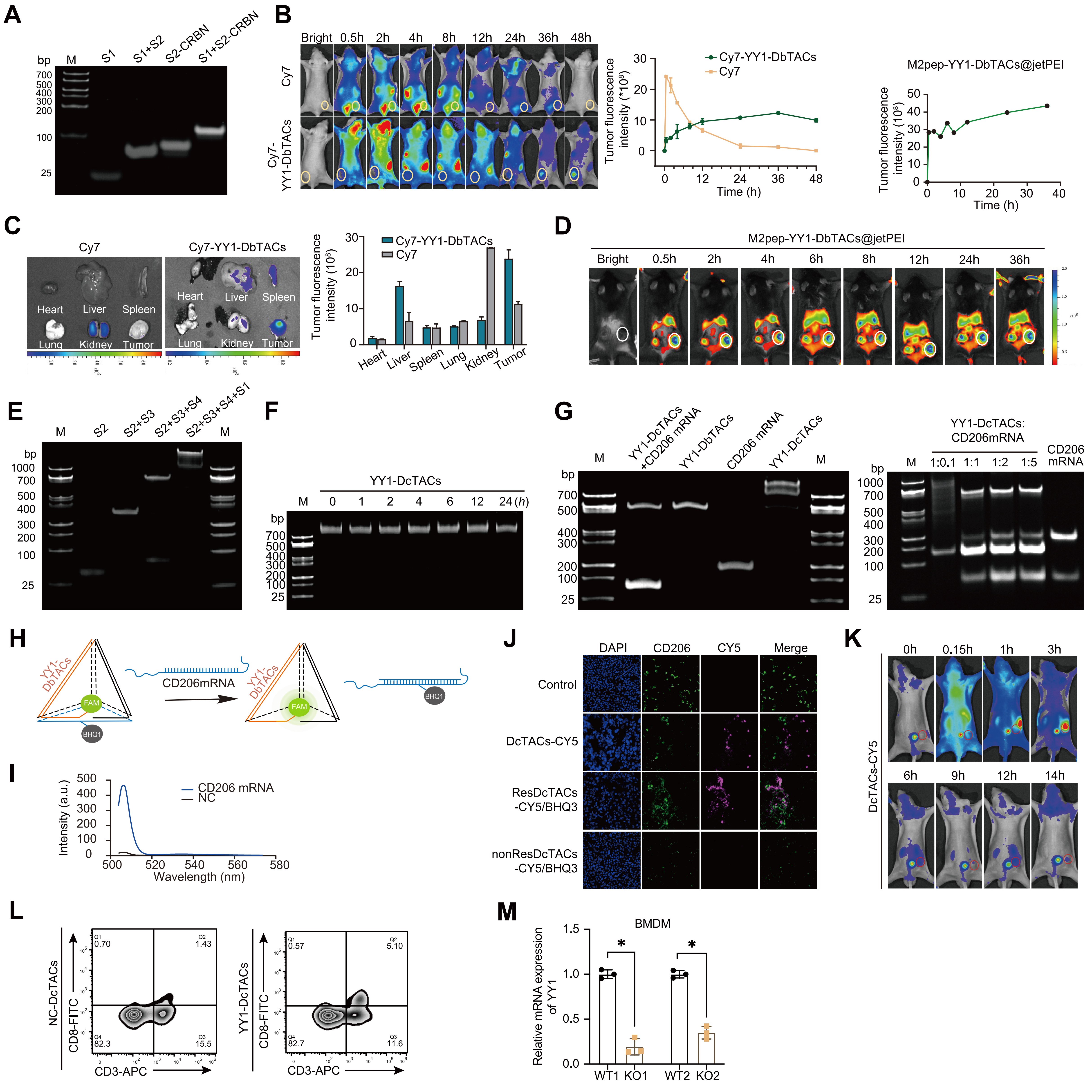
